# A second unveiling: haplotig masking of the eastern oyster genome improves population-level inference

**DOI:** 10.1101/2022.08.29.505626

**Authors:** Jonathan B. Puritz, Ximing Guo, Matt Hare, Yan He, LaDeana Hillier, Shubo Jin, Ming Liu, Katie Lotterhos, Pat Minx, Tejashree Modak, Dina Proestou, Edward S. Rice, Chad Tomlinson, Wes Warren, Erin Witkop, Honggang Zhao, Marta Gomez-Chiarri

**Affiliations:** University of Rhode Island, Department of Biological Sciences, Kingston, RI, USA; Haskin Shellfish Research Laboratory, Department of Marine and Coastal Sciences, Rutgers University, Port Norris, NJ, USA; Cornell University, Department of Natural Resources and the Environment, Ithaca, NY, USA; Department of Genome Sciences, University of Washington, Seattle, WA, 98195; Northeastern University, Department of Marine and Environmental Sciences, Nahant, MA, USA; University of Rhode Island, Department of Fisheries, Animal and Veterinary Sciences, Kingston, RI, USA; USDA Agricultural Research Service, National Cold Water Marine Aquaculture Center, Kingston, RI, USA; Bond Life Sciences Center, University of Missouri, Columbia, MO 6521; McDonnell Genome Institute, Washington University School of Medicine, St Louis, MO, USA; University of Missouri, Departments of Animal Sciences and Surgery, Institute for Informatics and Data Sciences, Bond Life Sciences Center, Columbia, MO 65211

**Keywords:** Genome Assembly, haplotig, population genomics, genetic diversity, genome curation

## Abstract

Genome assembly can be challenging for species that are characterized by high amounts of polymorphism, heterozygosity, and large effective population sizes. High levels of heterozygosity can result in genome mis-assemblies and a larger than expected genome size due to the haplotig versions of a single locus being assembled as separate loci. Here, we describe the first chromosome-level genome for the eastern oyster, *Crassostrea virginica*. Publicly released and annotated in 2017, the assembly has a scaffold N50 of 54 mb and is over 97.3% complete based on BUSCO analysis. The genome assembly for the eastern oyster is a critical resource for foundational research into molluscan adaptation to a changing environment and for selective breeding for the aquaculture industry. Subsequent resequencing data suggested the presence of haplotigs in the original assembly, and we developed a *post hoc* method to break up chimeric contigs and mask haplotigs in published heterozygous genomes and evaluated improvements to the accuracy of downstream analysis. Masking haplotigs had a large impact on SNP discovery and estimates of nucleotide diversity and had more subtle and nuanced effects on estimates of heterozygosity, population structure analysis, and outlier detection. We show that haplotig-masking can be a powerful tool for improving genomic inference, and we present an open, reproducible resource for the masking of haplotigs in any published genome.

## 1 Introduction

A highly contiguous and annotated genome assembly can be an invaluable tool for population genomic inference (Ellegren, 2014) enabling both genome-scale and reduced representation methods for sequencing populations (Ekblom & Galindo, 2010; Fonseca et al., 2016; Matz, 2017). However, even as the cost of second and third generation sequencing continues to drop (van Dijk, Auger, Jaszczyszyn, & Thermes, 2014), many challenges remain for assembling references for non-model organisms (Roach, Schmidt, & Borneman, 2018; Solares, Tao, Long, & Gaut, 2021). One of the greatest hurdles to accurate genome assembly is heterozygosity (Kajitani et al., 2014; Safonova, Bankevich, & Pevzner, 2015; Vinson et al., 2005), a hallmark of many wild plant and animal species, including most insects and marine invertebrates.

One assembly artifact created by heterozygosity is the “haplotig”. Haplotigs arise when algorithms misinterpret divergent allelic haplotypes in heterozygous genomic regions and assemble them as separate (duplicated) loci, with both copies incorporated into the primary assembly (Roach et al., 2018). For a genomic region split into haplotigs in the assembled reference, read mapping software will randomly chose which haplotig to map a read to (i.e., half the reads map to one haplotig, and the other half of reads map to the other haplotig). This issue with read mapping means that haplotigs could present a major problem for population genomic analyses of any highly polymorphic species. The splitting of a diplotig into haplotigs might limit SNP discovery and reduce estimates of genetic diversity, as allelic reads could potentially map to different parts of the assembly, reducing coverage (needed for genotyping), allelic representation within reads for a specific locus, and at the worst extreme producing two homozygous genotypes for what in reality would be a single heterozygous SNP. If the haplotigs were associated with populations (one allele more common in one subpopulation than another), this could introduce errors into statistics like population estimates of observed and expected heterozygosity as well as *F_ST_*. Haplotigs could also affect estimates of copy number variation and other structural variants by inherently making extra copies of loci within the genome. Moreover, haplotig detection is still a developing field in bioinformatics with most software published within the last four years (Pryszcz & Gabaldón, 2016; Roach et al., 2018; Solares et al., 2021). As a result, many published genome assemblies likely have some amount of haplotigs incorporated into the assembly, and the extent to which haplotig artifacts impact population genomic analysis has not been well studied.

The eastern oyster, *Crassostrea virginica*, is an excellent species to understand the impacts of haplotigs on population genomic inference. Eastern oysters create three dimensional reef structures that provide multiple ecosystem services such as providing habitat for nearshore fishes, water quality improvements, and protection from storm waves and erosion; the value of oyster reef ecosystem services can be as much as $99,000 per hectare per year (Grabowski, Conrad, & James, 2012). Additionally, the eastern oyster supports a commercial fishing and aquaculture industry valued at $100 million with several regional selective breeding programs (Allen, Small, & Kube, 2021; Gómez-Chiarri, Warren, Guo, & Proestou, 2015; Guo, 2021). Eastern oysters are highly fecund, protandrous hermaphrodites with external fertilization and a planktonic larval stage (Thompson, Guo, & Harrison, 1996), are distributed from New Brunswick, Canada, to Yucatan, Mexico, and are genetically diverse (Buroker, 1983; Hoover, Cindi A., and Patrick M. Gaffney, 2005; Karl & Avise, 1992; Reeb & Avise, 1990; Varney, Galindo-Sánchez, Cruz, & Gaffney, 2009).

The genome assembly of the eastern oyster is scaffolded onto the 10 known chromosomes for this species and was first released on NCBI in 2017. Initial results from a resequencing project indicated that there may be haplotig sequences in the genome, so we developed a protocol to infer haplotigs within the assembly. In this paper, we describe the assembly and annotation of the eastern oyster genome. We then describe a *post hoc* protocol to detect and mask haplotigs in a genome assembly and apply that to the eastern oyster genome. Lastly, we compare the haplotig-masked genome to the original genome to understand the effects of haplotigs on read mapping, SNP discovery, and estimates of common statistics in population level inference. We show that haplotig-masking can be a powerful tool for improving population genomic analysis without having to reassemble a genome.

## 2 Materials and Methods

### 2.1 Original Genome Sequencing and Assembly

#### 2.1.1 Sequenced individual

An inbred gynogenetic (with DNA from the mother only) oyster (RU13XGHG1#28) was used for whole genome sequencing to reduce problems associated with high polymorphism. Inbred gynogenetic oysters were produced by meiotic gynogenesis (Guo, Hershberger, Cooper, & Chew, 1993) from an oyster line (NEH^®^) selectively bred for 12 generations. Samples of tissue from parents and progeny were flash frozen in liquid nitrogen and stored at -80° C. Levels of inbreeding in the gynogenetic progeny as compared to the parents and wild oysters from Delaware Bay were determined using a panel of 15 microsatellites (Wang, Wang, Wang, & Guo, 2010), and a confirmed inbred female oyster from the gynogenetic progeny was selected for sequencing.

#### 2.1.2 Sequencing and assembly

High quality genomic DNA from the gynogenetic individual was isolated using the Trizol method from a pool of two tissues (gill, mantle), and DNA quality was assessed through Nanodrop, Qubit, and TapeStation analysis. All sequences were generated on a PacBio RSII instrument with P6-C4 sequencing chemistry. De novo assembly used PacBio subreads (>8 kb) with the standard FALCON v0.5.0 (Chin et al., 2013) method (parameter setting: max_diff 120, max_cov 120, min_cov 3, min_seed_length 9Kb) and assembled contigs were error-corrected with Quiver (Chin et al., 2013). Error corrected haplotype-specific contigs were first linked together into scaffolds using SSPACE-LongRead (Boetzer & Pirovano, 2014). Scaffolds were then linked together into megascaffolds using a high-order chromatin contact map (HiC) between chromosomes (Bickhart et al., 2017). Adductor muscle tissue from a reference related individual was used to generate HiC libraries at Phase Genomics (Seattle, WA) for this purpose. The libraries were sequenced using paired end sequencing on a HiSeq X Illumina instrument and reads (100bp) were aligned to the error-corrected contigs within scaffolds using BWA V0.7.16 (Li & Durbin, 2010) with strict parameters (-N 0; no hits from discordant pairs) to prevent mismatches and nonspecific alignments. Only read pairs that aligned to different contigs were used for scaffolding. The Proximo Hi-C pipeline performed chromosome clustering and contig orientation as described previously (Bickhart et al., 2017). At this stage all pseudochromosome sequences were polished with Pilon (Walker et al., 2014) using ∼40x coverage of Illumina data from the same reference individual.

### 2.2 Original Assembly curation

#### 2.2.1 Annotation

To develop transcript resources, we extracted total RNA from muscle, digestive, gill, and mantle tissues, and pools of larvae using a RNeasy kit (Qiagen) according to the manufacturers’ protocol. Total RNA for quality was assessed on the Agilent Fragment Analyzer, then enriched for poly(A)+ RNA using the MicroPolyA Purist kit (Ambion, Carlsbad, CA). We used ScripSeq (Epicentre, Madison, WI) to generate strand-specific cDNA that was sequenced on the Illumina Hiseq4000 platform as 100 base paired-end reads (insert size of 400bp). All RNAseq tissue data (150 million reads) were assembled with Trinity version 2.1.0 (Grabherr et al., 2011). The open reading frames (ORFs) were extracted from the complete transcriptome assembly (Trinity.fasta) using TransDecoder and LongOrfs modules (https://github.com/TransDecoder/TransDecoder/releases/tag/TransDecoder-v5.5.0). The 819Mb primary de novo assembly was used as input to align Trinity assembled transcripts using BLAT (Bhagwat, Young, & Robison, 2012; James Kent, 2002). Top hits were parsed with internal scripts by requiring there only be one best hit (-total_hits 1) with subsequent scaffolds showing multiple transcript alignments manually inspected for possible redundancy using assembly self-alignment data.

For assembly self-alignment we first fragmented the assembly in silico into 1 kbp segments and aligned against itself using BLASTZ (Schwartz et al., 2003). These alignments are scored against a repeat masked reference sequence using RepeatModeler (http://www.repeatmasker.org/RepeatModeler) output that is suitable for RepeatMasker application (N. Chen, 2004). After several alignment criteria were evaluated, we used 97% identity and 80% coverage with each scaffold to remove redundant contigs. For gene completeness, all assembled transcripts were aligned against the pseudochromosomes using BLAT requiring there only be one best hit (-total_hits 1) and parsed by varied alignment length thresholds of 95, 75 and 25% at a 90% sequence identity cutoff. Cumulative representation was summed for all transcript varied length alignments. The NCBI pipeline used for the gene annotation of *C. virginica* genome followed methods detailed in Pruitt et al. (Pruitt, Tatusova, Brown, & Maglott, 2012). Lastly, a genetic linkage map with 4006 RAD-seq markers was constructed with data from 115 progeny from an F2 family (He, 2012) using JoinMap 4.0 (Van Ooijen, 2006) and used to assess the integrity of the assembly. Accuracy of sequence placement was assessed using genetic marker sequence alignments against the *C. virginica* linkage map as defined by Chromonomer (Catchen, Amores, & Bassham, 2020) ordering that also uses the genetic linkage map as input.

### 2.3 Haplotig Detection and Masking

The original curated and annotated assembly (described above) was deposited in NCBI (RefSeq Accession: 4991078) in September of 2017 for widespread usage by several stakeholders. Afterwards, a resequencing project began and during preliminary data analysis in late 2019, a pattern in coverage across the genome began to indicate that there may be haplotig sequence contained within the genome assembly. To investigate, the genomic Illumina reads from the original genome individual (used for genome polishing) were mapped to the original assembly using BWA (Li & Durbin, 2010). Coverage was averaged across 10kb windows using samtools (Li et al., 2009) and plotted as a histogram using R (R Development Core Team, 2008). The bimodal distribution of coverage with a peak at expected coverage (diploid; 64X) and half the expected coverage (haploid; 32X) confirming the presence of haplotigs in the original assembly. Because the genome had already been publicly released, and widely used, we chose to mask haplotigs while preserving the original genome coordinates, enhancing compatibility with previous and ongoing research.

Files, scripts, and an RMarkdown file to reproduce the entire haplotig detection and masking process can be found at (https://github.com/The-Eastern-Oyster-Genome-Project/2022_Eastern_Oyster_Haplotig_Masked_Genome/tree/main/Haplotig_Masking; with an archived release of the code for this submission accessible at <u>DOI: 10.5281/zenodo.7448959</u>). To detect haplotigs, the scaffolded assembly was broken into the original set of assembled contigs by converting the NCBI annotation file (GCF_002022765.2_C_virginica-3.0_genomic_gaps.txt.gz) to a bed file and using BEDTools (Quinlan & Hall, 2010) to “subtract” gaps from the fasta file, generating a set of 669 contigs. To test for chimeric contigs, the genomic Illumina reads from the original genome individual were then mapped to these 669 contigs using minimap2 (Li, 2018, 2021) and processed with samtools (Li et al., 2009) and Picard Tools (Institute, 2016) to remove poorly mapping reads and optical and PCR duplicates. BEDTools (Quinlan & Hall, 2010) was then used to calculate mean coverage across sliding 100 kb windows with 50 kb of overlap.

We used coverage windows to break up chimeric contigs by looking for contigs that had large windows (>150kb) that shifted from diploid coverage levels to haploid coverage levels. We used BEDTools (Quinlan & Hall, 2010) to calculate these interval changes to break potentially chimeric contigs into multiple smaller contigs, preserving all basepairs. The coverage across sliding windows was used to split potentially chimeric contigs by initially dividing windows into high (> 40 mean coverage: all-high-intervals) and low (<40 mean coverage: all-low-intervals) with remaining windows merged. A coverage level of 40 was chosen based on the distribution of coverage which had a bimodal peak of 32 and 64; 40 represented the tail end of the “haplotig” peak. Remaining windows smaller than 150kb were also extracted from each of the two coverage sets, as potential chimeras.

Our logic for breaking contigs, focused on length and coverage of the windows. The all-high-intervals likely are all diplotigs given the coverage, and the low coverage windows smaller than 150kb are likely to be haplotigs that are nested within chimeric contigs. We combined this interval set, representing high-confidence diplotigs and high-confidence chimeric haplotigs (Subset 1). The other two interval sets, the low coverage intervals larger than 150kb and the high coverage intervals smaller than 150kb are merged, representing high confidence haplotig intervals and high confidence diplotigs that are nested within chimeric contigs (Subset 2). Subtracting the high confidence diplotig intervals and the likely nested haplotig intervals (Subset 1) from the high-confidence haplotigs and likely chimeric diplotigs (Subset 2), breaks any potential chimeras. The sequence in the broken contigs is recovered by extracting sequences from the original reference using Subet 1 and the result of Subset 1 - Subset 2, producing a set of contigs broken at chimeric points.

Minimap2 (Li, 2018, 2021) was then used to map the Illumina reads to the broken contigs, and samtools (Li et al., 2009) converted alignments to a BAM file. The program Purge Haplotigs (Roach et al., 2018) was used to then detect and remove haplotigs from the manual reference contigs. The “coverage” step of the program was run with 10 as the “low cutoff” (-l), 56 as the “midpoint (-m) and 120 was the “high cutoff” (-h). The ‘purge’ and ‘clip’ steps were run with default parameters. The mitochondrial genome was flagged in the initial purge step as an artifact due to its high coverage. It was searched for and removed from the list of artifact sequences. We then combined the haplotigs from the clip step, and the initial artifacts to create a “total haplotig” file. We mapped this file back to the original reference, using minimap2, to identify haplotigs, and used BEDTools (Quinlan & Hall, 2010) to convert the mappings to a haplotig bed file. This haplotig bed file was used to mask the reference genome using BEDTools to produce the final haplotig-masked genome.

### 2.4 Genome comparison

All code, scripts, and files needed to reproduce the comparative analysis can be found in the https://github.com/The-Eastern-Oyster-Genome-Project/2022_Eastern_Oyster_Haplotig_Masked_Genome repository in the folder “Comparative_Analysis”

#### 2.4.1 Coverage

To assess the impacts of haplotig masking, Illumina reads from the original genome individual were mapped to both genome versions. A modified version of the dDocent pipeline (Puritz, Hollenbeck, & Gold, 2014) was used to run bwa (Li & Durbin, 2010) to map reads to the genome. Duplicate reads were identified with the “MarkDuplicates” function of Picard (Institute, 2016). Samtools (Li et al., 2009) was then used to create individual bam files based on a set of filtering criteria: “total reads”-bam file contains all non-duplicate primary alignments, “multimapping reads”-bam file contains only reads that mapped to more than one location in the genome, and “filtered reads”-bam file contains only has mapping with a quality score above 10 and no hard or soft clipping above 80bp. BEDTools (Quinlan & Hall, 2010) was then used to calculate the average coverage over 10kb windows across the genome. Coverage was plotted as a histogram using the ggplot2 package (Wickham, 2016) in R (R Development Core Team, 2008). Coverage was also plotted along individual chromosomes.

#### 2.4.2 Completeness of genome

Genome completeness was assessed for both versions using Benchmarking Universal Single-Copy Orthologs (BUSCO) version 5.4.3 (Seppey, Manni, & Zdobnov, 2019; Simão, Waterhouse, Ioannidis, Kriventseva, & Zdobnov, 2015) and the Mollusca ortholog database version 10, containing 5295 single-copy orthologs. For comparison, the two newest chromosome level assemblies for *Crassostrea gigas* were downloaded from NCBI (GCA_011032805.1, GCA_902806645.1) and assessed using the same BUSCO version and database.

### 2.5 Population level inference

#### 2.5.1 Resequencing data

Ninety adult wild and farmed eastern oysters were collected in the fall of 2017 from multiple water bodies across the United States of America including the Gulf of Maine, the Delaware Bay, the Chesapeake Bay, and the northern Gulf of Mexico near Louisiana. Samples were also included from multiple selected oyster lines, for a total of eight wild localities and five selected lines. Individuals were sequenced on an Illumina HiSeq X PE 150 bp platform to 15-20X coverage. Twelve samples were included in sequencing and variant calling from known inbred experimental lines and populations as part of a different research project. These samples were included for mapping statistics and SNP counts, but were not used for any population level analyses, leaving a total of 78 individuals. Full details on sample source, and collection, processing, and sequencing methods can be found in <u>(Puritz et al., 2022)</u>.

#### 2.5.2 Nucleotide Variant Calling

Raw sequencing reads were processed with a modified version of the dDocent pipeline (Puritz et al., 2014). First, reads were trimmed for low quality bases and adapter sequences using the program fastp (S. Chen, Zhou, Chen, & Gu, 2018). Trimmed reads were mapped to both genome versions using bwa (Li & Durbin, 2010) with mismatch and gap-opening parameters (-B 3 -O 5). Picard (Institute, 2016) was used to mark duplicate reads, and subsequent BAM files were filtered with samtools (Li et al., 2009) to remove low quality mappings, secondary alignments, and PCR duplicates. The program freebayes (Erik Garrison & Marth, 2012) was used to genotype small nucleotide variants (SNPs, InDels, small complex events).

Bcftools (Danecek et al., 2021) and vcftools (Danecek et al., 2011) were used in combination to filter raw variants. Variants were filtered based on allelic balance at heterozygous loci (between 0.1 and 0.9) and quality to depth ratio of greater than 0.1. Variants were then filtered based on mean-depth, excluding all loci above the 95th percentile. Vcflib (E. Garrison, 2016) was then used to decompose variants into SNPs and InDels. Lastly, SNPs were filtered to allow for no missing data and only biallelic SNPs, and then variants separated into two sets of variants, one with a minor allele frequency (MAF) of 1% and the other with a MAF of 5%.

#### 2.5.3 Structural variant calling

We used the program Delly (Rausch et al., 2012) following the “germline sv calling” (https://github.com/dellytools/delly#germline-sv-calling) pipeline to identify candidate structural variants (SVs), including deletions, insertions, duplications and inversions. SVs were filtered using Delly with the “germline” filter. BCFtools (Danecek et al., 2021) was used to convert inversion SVs to a bed file and then switch “LowQual” genotypes to missing, and SVs were filtered to a subset with no missing data. Using this filtered SV subset, read based copy number (VCF Format ID = RDCN) for insertions, deletions, and duplications were extracted to a tab delimited list for both genome versions.

#### 2.5.4 Identification of newly diplotig regions in haplotig-masked genome

When a haplotig is effectively masked, the remaining haplotig should become a diplotig. We identified these regions to see if changes in population inference were more pronounced in these regions relative to the rest of the genome. BEDTools (Quinlan & Hall, 2010) was then used to calculate the average coverage over 10kb windows across both genome versions. New diplotig regions were identified as any 10kb window that increased in coverage in the haplotig-masked genome by greater than 1.5 times the coverage in the original genome version.

#### 2.5.5 Nucleotide diversity

Nucleotide diversity (π) was calculated across 10kb windows of both genomes using VCFtools (Danecek et al., 2011) with the SNP dataset with greater than 1% minor allele frequency. Differences between the original and haplotig-masked nucleotide diversity estimates were tested using a t-test. π was visualized across genomic windows for both genome versions, as was the difference between the two estimates. Differences between estimates from the two genome versions were also visualized and tested across new diplotig regions.

#### 2.5.6 Heterozygosity

Vcflib (E. Garrison, 2016) was used to calculate per-site values of SNP heterozygosity for the SNP dataset with only biallelic SNP with greater than 1% minor allele frequency. Per-site values were averaged across 10kb windows using the program BEDTools (Quinlan & Hall, 2010) for both versions of the genome. Differences between original and haplotig-masked heterozygosity estimates were tested and visualized in the same way as nucleotide diversity, across the whole genome and only in new diplotig regions.

#### 2.5.7 Global *F_ST_* and outlier detection

The program OutFLANK (Whitlock & Lotterhos, 2015) was used to calculate global *F_ST_* for biallelic SNPs with a minor allele frequency greater than 5%. Outliers were inferred relative to a null *F_ST_* distribution based on trimmed SNP datasets with heterozygosity greater than 0.1 and using a random set of 50,000 independent SNPs derived from snp_autoSVD in bigsnpr (Privé, Aschard, Ziyatdinov, & Blum, 2018) using the settings (min.mac = 7, size=10). OutFLANK calculated q-values for outlier scores for all SNP loci with heterozygosity above 0.1. A false discovery rate of 0.05 was used to designate significance based on q-values.

The full set of oyster individuals was found to have significant population structure between the Gulf of Mexico and Atlantic wild populations, as well as among selected lines and wild populations (pairwise *F_ST_* ∼ 0.1-0.5; Puritz et al. 2022). To examine patterns in *F_ST_* in a lower structure dataset (pairwise *F_ST_* ∼ 0.01-; Puritz et al. 2022), populations were subset to wild populations only from the Atlantic coast of the USA (LSS-6 populations, 36 individuals). For the LSS, snp_autoSVD was run with the settings (min.mac =4, size=10) to account for the smaller number of individuals but OutFLANK run options remained the same. Differences between the original and haplotig-masked *F_ST_* values were tested using a t-test across both the full data and the LSS subset. *F_ST_*values were also visualized as Manhattan plots for: the original genome, the haplotig masked genome, and the difference between the estimates. For visualization, *F_ST_* values were averaged across 10kbp windows. If a single outlier SNP was detected in a 10kb window, the entire window was visualized as an outlier. Lastly, *F_ST_* values were also tested for differences and examined across new diplotig regions in all datasets.

#### 2.5.8 Copy number differentiation

Copy number variants (CNVs) were filtered for a minor allele frequency greater than 5%. Differentiation at CNVs was calculated using the *V_ST_*statistic (Redon et al., 2006) as implemented in (Steenwyk, Soghigian, Perfect, & Gibbons, 2016) and plotted across the genome. The difference in *V_ST_* values produced from the different genome versions across various data subsets was tested using a t-test and visualized across chromosomes in Manhattan plots similar to *F_ST_*values.

## 3 Results

### 3.1 Original assembly

#### 3.1.1 Sequenced individual

The individual sequenced was from a family produced by gynogenesis of an already inbred female oyster. Genotyping with a panel of 11-15 microsatellite loci showed that this family experienced an approximately 55.4% reduction in the heterozygosity compared with their parents. The average heterozygosity in the gynogenetic progeny was 0.115, compared with 0.642 in wild Delaware Bay oysters (Supplemental Table 1).

#### 3.1.2 Sequencing and assembly

We sequenced and assembled a reference genome for the eastern oyster using high-coverage paired-end libraries. We sequenced 11,116,776 PacBio reads (122.7 GB) resulting in 87x coverage. We also sequenced 138,800,932 paired-end Illumina reads that were used for polishing (and later genome assessment). We also generated over 690 million paired end reads for RNA transcript assembly and assembly annotation. All sequencing reads used for the assembly and curation can be found on NCBI with accession numbers found in Supplemental Table 2.

Our initial contig assembly of 819 Mb, was much larger than the genome size of 578Mb estimated by flow cytometry (Guo Lab, unpublished), and the Pacific oyster, *Crassostrea gigas,* assemblies of 647 and 586 Mb. This led us to utilize a strategy of genome self-alignment and duplicative transcript mapping that identified 135 Mb of heterozygous loci to remove. In the first assembly of the Pacific oyster polymorphic assembled loci were also removed in a similar way (Zhang et al 2012). Our final assembly consisted of 684 Mb in 669 contigs of N50 contig and scaffold length 1.97 and 54 Mb, respectively (Table 1). Most sequences (>99%) were scaffolded into the known number of 10 chromosomes using HiC and genetic linkage mapping data. The eastern oyster assembly represents a high level of contiguity (Table 2; Supplemental Table 3).

**Table 1.**
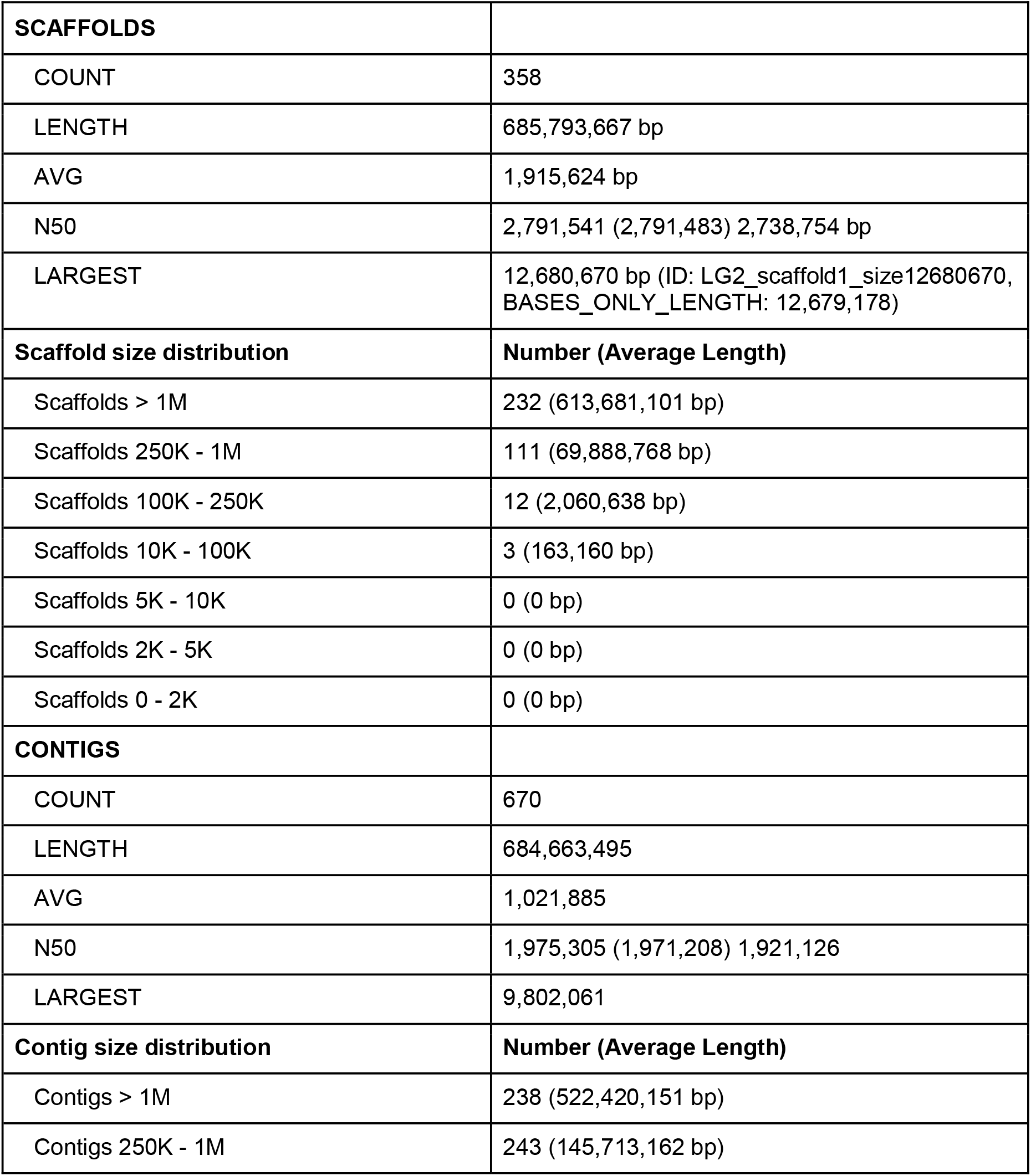
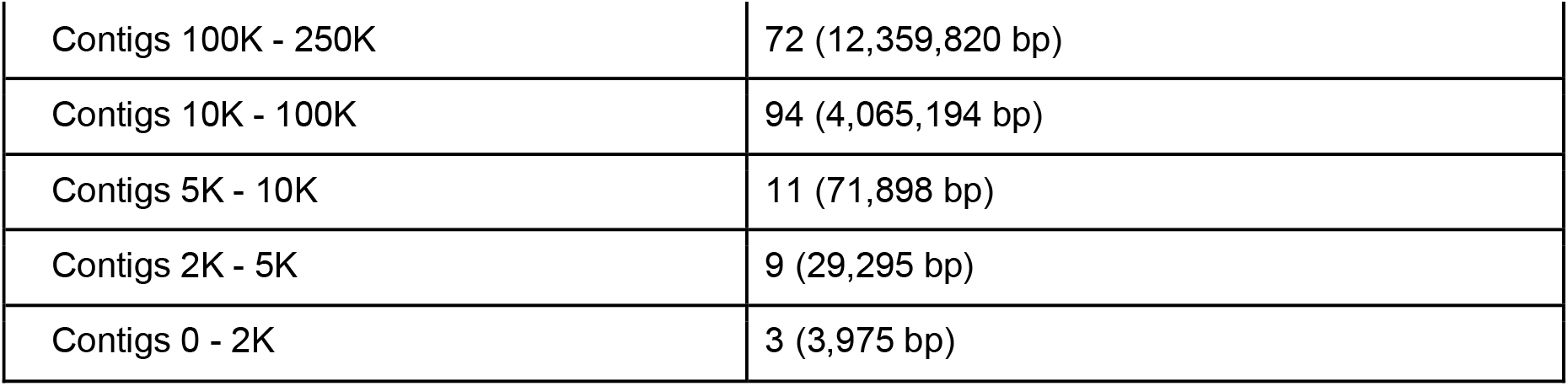
Assembly statistics:

|  |  |
| --- | --- |
| **SCAFFOLDS** |  |
| COUNT | 358 |
| LENGTH | 685,793,667 bp |
| AVG | 1,915,624 bp |
| N50 | 2,791,541 (2,791,483) 2,738,754 bp |
| LARGEST | 12,680,670 bp (ID: LG2_scaffold1_size12680670, BASES_ONLY_LENGTH: 12,679,178) |
| **Scaffold size distribution** | **Number (Average Length)** |
| Scaffolds > 1M | 232 (613,681,101 bp) |
| Scaffolds 250K - 1M | 111 (69,888,768 bp) |
| Scaffolds 100K - 250K | 12 (2,060,638 bp) |
| Scaffolds 10K - 100K | 3 (163,160 bp) |
| Scaffolds 5K - 10K | 0 (0 bp) |
| Scaffolds 2K - 5K | 0 (0 bp) |
| Scaffolds 0 - 2K | 0 (0 bp) |
| **CONTIGS** |  |
| COUNT | 670 |
| LENGTH | 684,663,495 |
| AVG | 1,021,885 |
| N50 | 1,975,305 (1,971,208) 1,921,126 |
| LARGEST | 9,802,061 |
| **Contig size distribution** | **Number (Average Length)** |
| Contigs > 1M | 238 (522,420,151 bp) |
| Contigs 250K - 1M | 243 (145,713,162 bp) |
| Contigs 100K - 250K | 72 (12,359,820 bp) |
| Contigs 10K - 100K | 94 (4,065,194 bp) |
| Contigs 5K - 10K | 11 (71,898 bp) |
| Contigs 2K - 5K | 9 (29,295 bp) |
| Contigs 0 - 2K | 3 (3,975 bp) |

**Table 2.** Comparison to published molluscan genomes. Table summarizes data from comparable molluscan genomes. For genomes where completeness was evaluated with BUSCO, the metazoan reference was used except for *C. virginica* which used the Molluscan reference, P. fucata martensii and Octopus minor where the reference was not specified. ^†^Evaluated by mapping short reads,sanger-sequenced BACs, and transcripts to the assembly ^‡^Only reported chromosome scaffolds. Unclear if there were unplaced scaffolds. ^§^Evaluated by mapping transcripts with predicted ORFs back to the genome ^¶^23,509 *<u>O. bimaculoides</u>* genes were covered at 90% of coding sequence by *O. vulgaris* reads. ^††^Evaluated with BUSCO and CEGMA but did not include actual completeness numbers; said that this assembly was less complete than previous versions, but most duplication issues were resolved

| Species | Publication | Individual sequenced | Genome Size (Mb) | Number of Chromosomes Assembled | Scaffold N50 | Scaffold # | Completeness | Genes | Repeat % |
| --- | --- | --- | --- | --- | --- | --- | --- | --- | --- |
| *Crassostrea virginica* | This study | Gynogen after 12 generations of inbreeding | Original- 685.7<br>Masked- 584.3 | 10 | 54,000,000 | 358 | Original- 97.4%<br>Masked-97.2% | 34,596 | 33% |
| *Lottia gigantea* | (Simakov et al., 2013) | Wild single male gonad | 359.5 | NA | 1870 | 4475 | NA | 23800 | 21% |
| *Crassostrea gigas* | (Zhang et al., 2012). | 4th gen inbred female | 559 | NA | 401,319 | 401 | 95-99%<sup>†</sup> | 28027 | 36% |
|  | (Qi et al., 2021) | One female from a farm | 586.8 | 10 | 60,957,391 | 10<sup>‡</sup> | 92.5%<br>(BUSCO) | 30078 | 57.2% |
|  | (Peñaloza et al., 2021) | Inbred aquaculture female | 647.8 | 10 | 58,462,999 | 236 | 95.6%<br>(BUSCO) | 30724 | 43% |
| *Mytilus coruscus* | (Yang et al., 2021) | Wild female | 1566.5 | 14 | 99542347 | 4434 | 89.4%<br>(BUSCO) | 37478 | 47% |
| *Octopus bimaculoides* | (Albertin et al., 2015). | Wild | 2371.5 | NA | 466,100 | 379,696 | 97%<sup>§</sup> | 33638 | 45% |
| *Octopus minor* | (Kim et al., 2018) | Unknown | 5090 | NA | 196941 | 41584 | 76.2%<br>(BUSCO) | 30,010 | 44.% |
| *Mytilus galloprovincialis* | (Murgarella et al., 2016) | one ind from Vigo, Spain | 1,600 | NA | 2,600 | 1.7M | 16%<br>(CEGMA) | 10891 | 36% |
| *Pinctada fucata martensii* | (Du et al., 2017) | 3rd generation selected individual | 990 | NA | 324,000 | 939 | 82.8%<br>(BUSCO) | 32973 | Not reported |
| *Saccostrea glomerata* | (Powell et al., 2018) | single female; 6th generation from selective breeding program | estimated: 784<br>assembly: 788 | NA | 804,200 | 10,017 | 87.2%<br>(BUSCO and CEGMA) | 29738 | 45%; |
| *Haliotis rufescens* | (Masonbrink et al., 2019) | one female and one male cultured, bred from CA | estimated=1800;<br>assembly =1498 | NA | 1,900,000 | 8400 | 95.1%<br>(BUSCO) | 57785; | 33% |
| *Octopus vulgaris* | (Zarrella et al., 2019) | wild caught adult male | estimated=2798<br>assembly=1780 | NA | 263,097 | 77,683 | 50%<br>(BUSCO) | Unknown<sup>¶</sup> | >50% |
| *Haliotis laevigata* | (Botwright et al., 2019) | one cultured female | estimated=1540;<br>assembly=1760 | NA | 86,805 | 63,588 | 86.6%<br>(BUSCO) | 55,164 | NA |
| *Mytilus galloprovincialis* | (Gerdol et al., 2020) | single female | assembly 1280 | NA | 207,640 | 10,577 | NA<sup>††</sup> | 60,338 | 43% |

### 3.2 Original Assembly Curation

Gene annotation using the automated NCBI pipeline predicted the presence of 34,596 protein coding genes and 4,230 non-coding (Supplemental Table 4). When contrasted to the Pacific oyster genome (Peñaloza et al., 2021), we found a high percentage (36-40% total interspersed) of repetitive elements with two independent methods (Supplemental Table 5). From assembled transcripts aligned to our eastern oyster assembly, we found 87% of 171,712 transcripts at a 95% length cutoff. We estimate 22% of the repeats could not be assigned a classification suggesting additional work is needed to define the composition of oyster sequence repeats. The assembly annotation can be found on NCBI (GCF_002022775.2).

Five of the assembled chromosomes (1, 2, 3, 4 and 8) were correctly aligned with linkage groups (LGs) of the genetic map indicating that they were correctly assembled (Supplemental Tables 6 and 7). Three chromosomes (5, 6, and 9) were aligned to more than one LG at different regions suggesting that they represent misassembled chromosomes. Two chromosomes (7 and 10) corresponded to parts of LGs indicating that they are chromosomal fragments. There were some minor discrepancies between the assembly and genetic map that need to be resolved with additional data. The assembled genome size of 684Mb was 18.3% longer than the genome size of 578Mb estimated by flow cytometry (Guo Lab, unpublished), suggesting that the assembly still contains some allelic redundancy. One of the regions is a 1.1 Mb segment that appeared twice on Chr 1 (47,598,449-48,729,575 and 49,357,009-50,465,997). The duplicated segments had identical gene content and gene order. Duplicated gene pairs in the two segments had identical exon-intron structures and 98-100% similarity in coding sequences but varied greatly in intron sizes. The duplicated gene pairs included two copies of *alternative oxidase*, a single-copy gene in most invertebrates, which were 98% identical in coding sequence but differed greatly in intron sizes, and PCR amplification of intron 6 indicated the two copies were allelic haplotigs and not true paralogous duplications (data not shown).

### 3.3 Haplotig detection and masking

Breaking up chimeric contigs based on sequencing coverage resulted in 1852 contigs (from the original 669) with an N50 of 885,077 bp. The program Purge Haplotigs (Roach et al., 2018) identified 963 haplotigs (partial and whole contigs). This resulted in 1171 primary contigs (non-haplotigs) totaling 578,183,332 bp with an N50 of 9,802,061. To retain compatibility with past studies and chromosome-level scaffolding, haplotigs were masked from the original assembly by substituting “Ns” for haplotig bases. The final haplotig-masked genome contained the same 684,741,128 bp of the original assembly with 100,438,362 bp masked and is archived at <u>DOI: 10.5281/zenodo.7448959</u>. Assessed initially using the original Illumina sequencing reads from the assembly, the masked version of the genome had a higher overall mean coverage (50.615X) compared to the original assembly (45.6445X) with a pronounced shift in 10kb intervals with an approximate diploid coverage peak (∼65X) relative to intervals with haploid levels (∼32.5X) of coverage (Figure 1). Looking at total read mappings, multi-mapping reads, and filtered read mappings across 10kb chromosomal intervals, masked intervals showed a clear dip in coverage relative to diplotig regions in the original genome while in the masked version of the genome shows increased total and filtered read mappings in several regions (new diplotigs) while also decreasing overall rates of multi-mapping reads (Figure S1).

**Figure 1.**
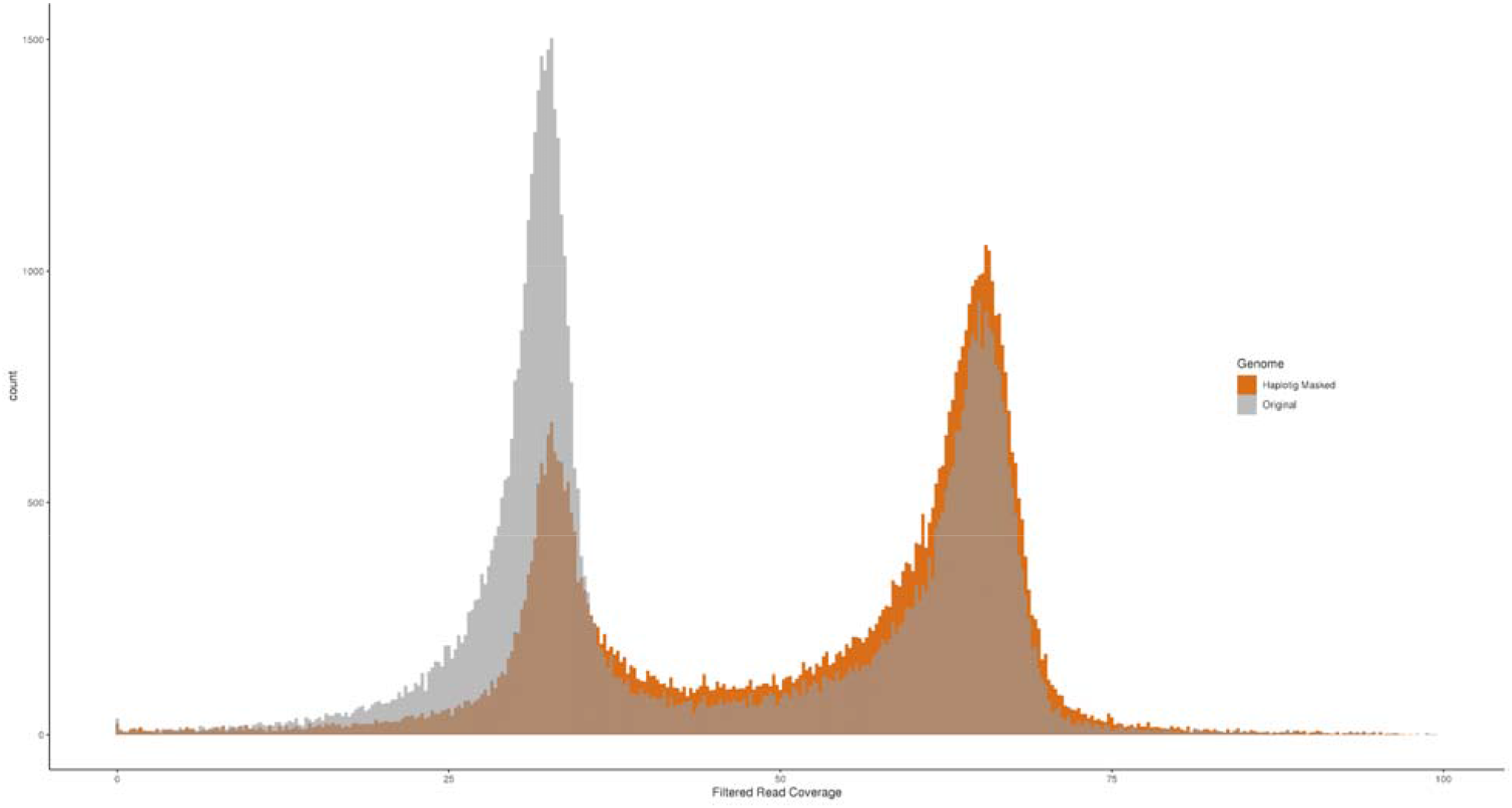
Histogram of read coverage across both genome versions. Paired-end Illumina reads used for polishing the original genome assembly were mapped back to the two genome versions. Filtered read coverage was averaged across 10kb windows and plotted as a histogram with bins colored by genome: gray for the original and orange for the haplotig-masked genome.

### 3.4 Haplotig masking increases read mappings and duplicate detections

We generated over 3,558,207,970 read pairs for the resequencing portion of this project with an average of 39,535,644 +/-1,018,131 read pairs per sample. On average, 97.14% +/− 2.49% of reads were retained after quality trimming and adapter removal. On average of 96.75% +/− 2.48% trimmed reads mapped to the original genome, compared with 97.01% +/− 2.49% to the haplotig-masked genome. For the original genome, 6.5% +/− 0.17% of mappings were marked as duplicates with 6.6% +/− 0.17% marked as duplicates for the haplotig-masked genome. Per sample statistics for sequencing, read mapping, and percent of the genome covered can be found in Supplemental Table S8.

### 3.5 Completeness of genome

Even though masking haplotigs removed over 100,000,000 bases from the original assembly, there was minimal impact on assembly completeness evaluated by BUSCO (Seppey et al., 2019; Simão et al., 2015). The original genome assembly had 5,158 complete (4,413 single copy, 745 duplicated), 34 fragmented, and 103 missing orthologs from the Mollusca-specific BUSCO database. In contrast, the haplotig-masked assembly version had 5,146 complete (5,034 single copy, 112 duplicated), 39 fragmented, and 110 missing orthologs (Figure S2). BUSCO assignment can be dependent on contig length, so BUSCO scores were also compared from non-scaffolded contigs. The original contigs had 5,156 complete (4,184 single copy, 972 duplicated), 34 fragmented, and 105 missing orthologs while the primary contigs (non-haplotigs) had 5,138 complete (5,003 single copy, 135 duplicated), 44 fragmented, and 113 missing orthologs. Using the Pacific Oyster, *C. gigas,* for comparison, the Qi et al. (2021) assembly of had 5,031 complete (4,836 single copy, 195 duplicated), 21 fragmented, and 243 missing orthologs and the Peñaloza et al. (2021) assembly had 5,198 complete (5,086 single copy, 112 duplicated), 26 fragmented, and 71 missing orthologs (Figure S3).

### 3.6 Haplotig contigs in assemblies reduce SNP discovery

Masking haplotigs increased the number of SNPs genotyped across all levels of filtering (Table 3). For the original assembly, 7,674,518 (3,580,098; MAF 5%) biallelic SNPs were kept after filtering compared to 12,149,052 (5,574,080; MAF 5%) for the haplotig-masked assembly. SNPs included in this 36.83% increase were found in representative proportions of genomic annotations (18.61% exonic, 53.54% intronic, 27.85% intergenic) with relatively even increases across categories (38.69% exonic, 35.74% intronic, 36.97% intergenic). The largest differences were in regions that switched from haploid to diploid coverage (new diplotigs) after masking (Figure 2). In new diplotigs (looking at all SNPs with a MAF > 0.01), the original assembly produced only 95,667 SNPs after filtering compared to 3,381,377 SNPs in the same regions of the haplotig-masked assembly with 3,306,188 (97.78%) exclusive only to the haplotig-masked genome. There were also 175,128 SNPs that were no longer present in the haplotig-masked genome with 98,589 of those SNPs found inside of haplotigs and 20,478 found inside new diplotigs. Out of the 8,735,612 SNPs that were called within both genome versions, genotypes had a mean 99.62% concordance rate, and this rate was lower in new diplotigs (98.97%) vs other regions of the genome (99.97%). Along with the number of SNPs genotyped, masking haplotigs had significant effects on the levels of inferred nucleotide diversity, heterozygosity, and *F_ST_*values across the genome.

**Figure 2.**
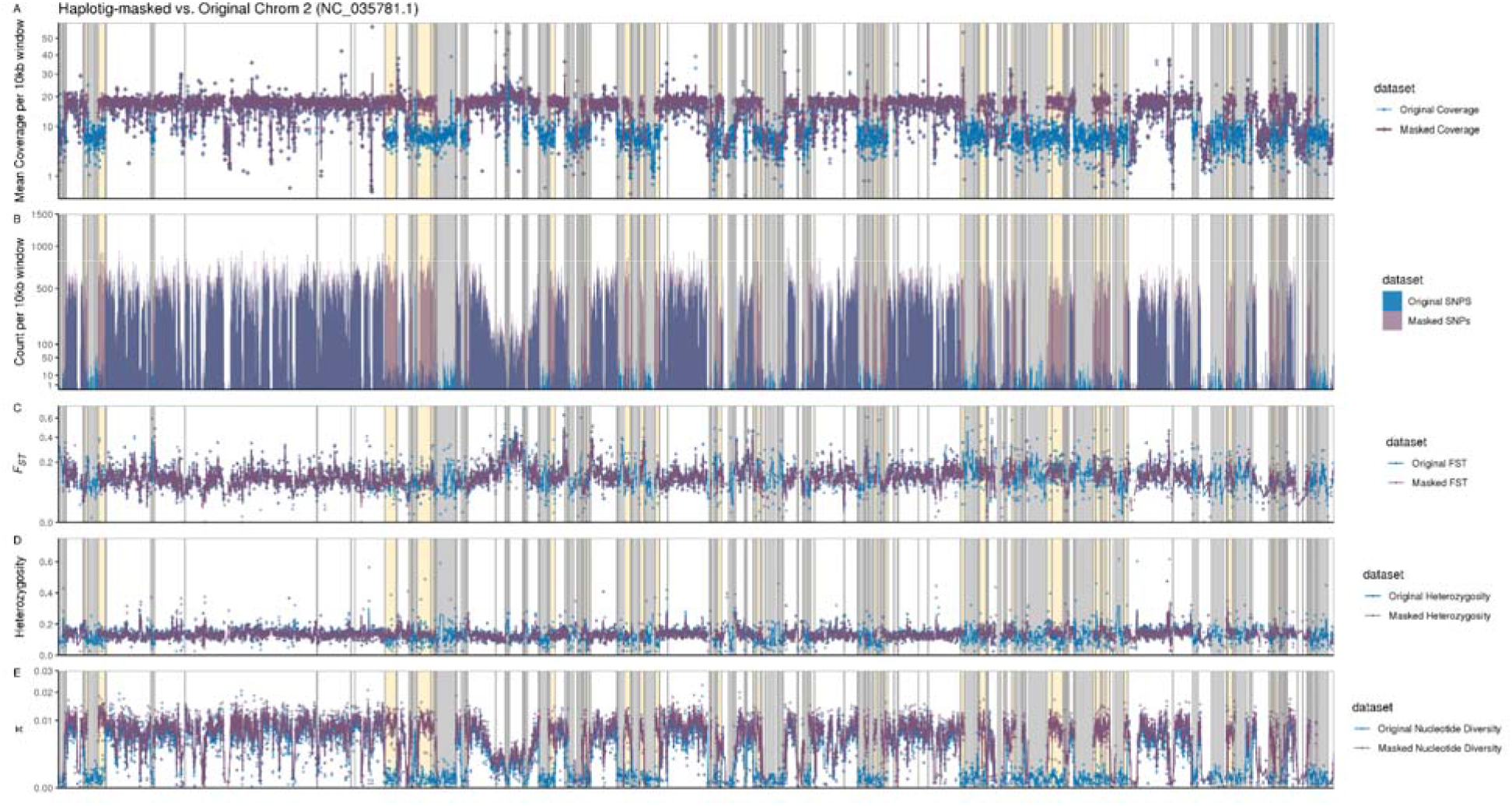
Comparison of coverage, SNPs, *F_ST_*, Heterozygosity and Nucleotide diversity across original and haplotig-masked assembly. Across both assemblies, coverage, the number of SNPs, *F_ST_*, heterozygosity, and nucleotide diversity were averaged across ten kilobase windows of Chromosome 2 (NC_03578.1). For coverage (Panel A), *F_ST_* (Panel C), heterozygosity (Panel D), and nucleotide diversity (Panel E), points are the values per 10 kb window with lines drawn as rolling 3-point averages, and the number of SNPs are plotted as an area plot. Areas along the chromosome shaded in gray were identified as haplotigs and therefore have no data for the haplotig-masked genome. Areas shaded in yellow are non-masked regions that showed a shift from haploid to diploid coverage levels after haplotig-masking. For all plots, blue is the original genome and purple is the haplotig-masked genome.

**Table 3.** SNP Results.

| Filtering Level | Original Genome | Haplotig-masked Genome |
| --- | --- | --- |
| Initial bioinformatic filtering | 45,427,924 | 52,971,541 |
| No missing data, MAF >0.01 | 8,910,740 | 14,103,332 |
| No missing data, MAF >0.01, 2 alleles | 7,674,518 | 12,149,052 |
| No missing data, MAF >0.05 | 4,299,397 | 6,699,719 |
| No missing data, MAF >0.05, 2 alleles | 3,580,098 | 5,574,080 |
| Low Structure Subset |  |  |
| No missing data, MAF >0.05, 2 alleles | 2,872,577 | 4,482,328 |

### 3.7 The presence of haplotigs greatly reduces estimates of nucleotide diversity

Across all calculated measures of genomic diversity and structure, nucleotide diversity (π) was significantly and drastically affected by the presence of haplotigs in the genome assembly (Figure 2; Figure 3). For comparison, values of π were averaged across 10kb windows, and for the original assembly the genome-wide average was 0.00382 +/− 1.93 x 10^-5^ compared to haplotig-masked 0.00587 +/− 2.20 x 10^-5^, and these two values differed significantly when evaluated with a t-test (*t = 70.2;* df=74574*, p =* 0). When individual window values are visualized across the whole genome (Figure 3) or across single chromosomes (Figure 2) clear drops in diversity line up with identified haplotig regions. There was also a clear increase in estimates of diversity in new diplotigs (Figure 4). Compared across new diplotigs only, the difference in nucleotide diversity was over an order of magnitude, with the original assembly average calculated to be 0.000321 +/− 7.6772 x 10^-6^ compared to 0.00720 +/− 4.68 x 10^-5^ for the haplotig-masked assembly. This difference was also significant when evaluated with a one-sided t-test (*t* = 145; df= 7794. *p* = 0).

**Figure 3.**
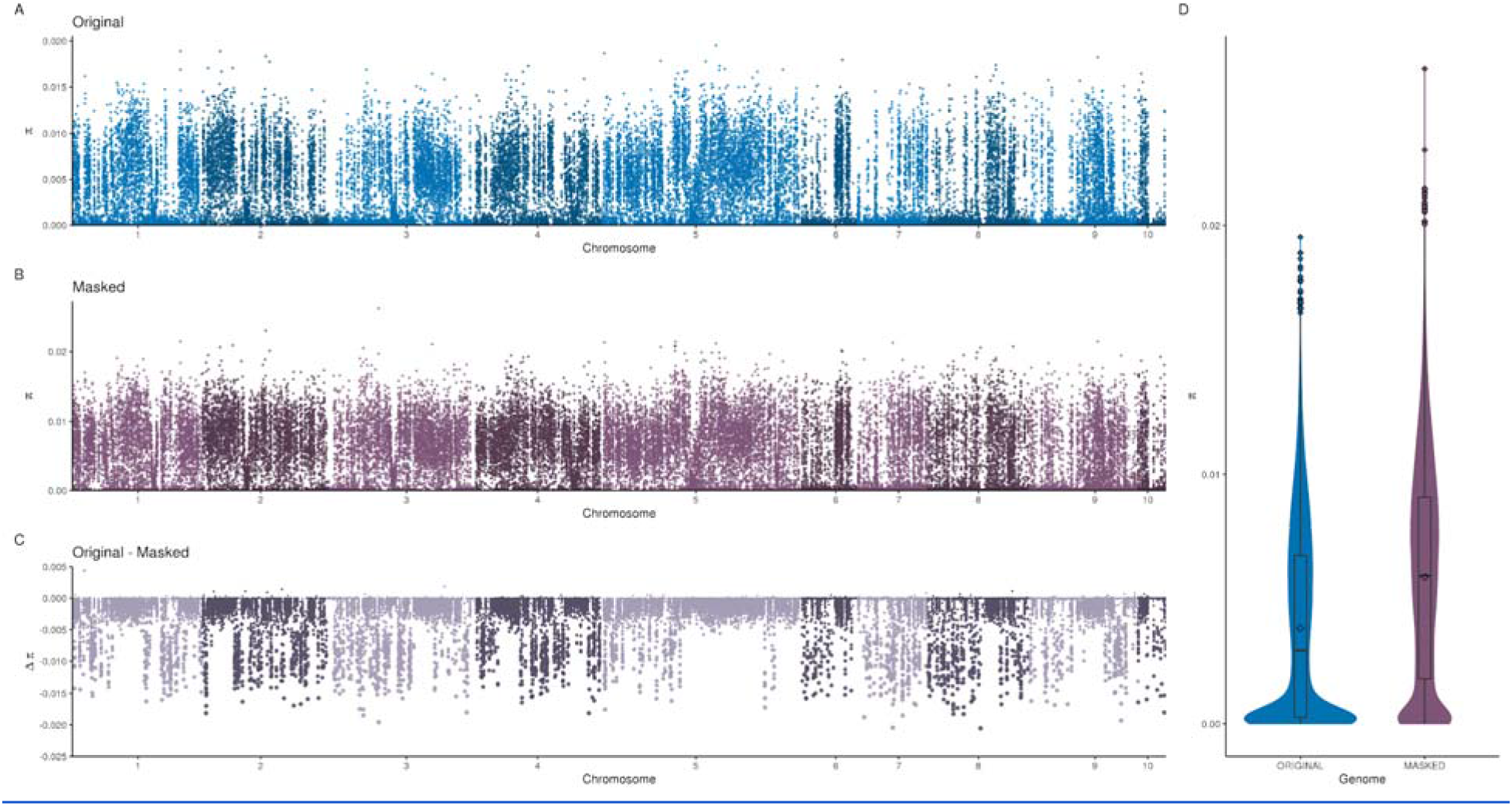
Comparison of nucleotide diversity across the original and haplotig-masked assembly. Panel (A) is values of π averaged across 10kb windows across the original genome. Panel (B) panel is values of π averaged across 10 kb windows across the haplotig-masked genome. Panel (C) is the difference between the original and the haplotig-masked values in 10kb windows across the entire genome. Panel (D) is a violin and boxplot of 10kb averaged values between the two genome versions. For all plots, blue is the original genome and purple is the haplotig-masked genome.

**Figure 4.**
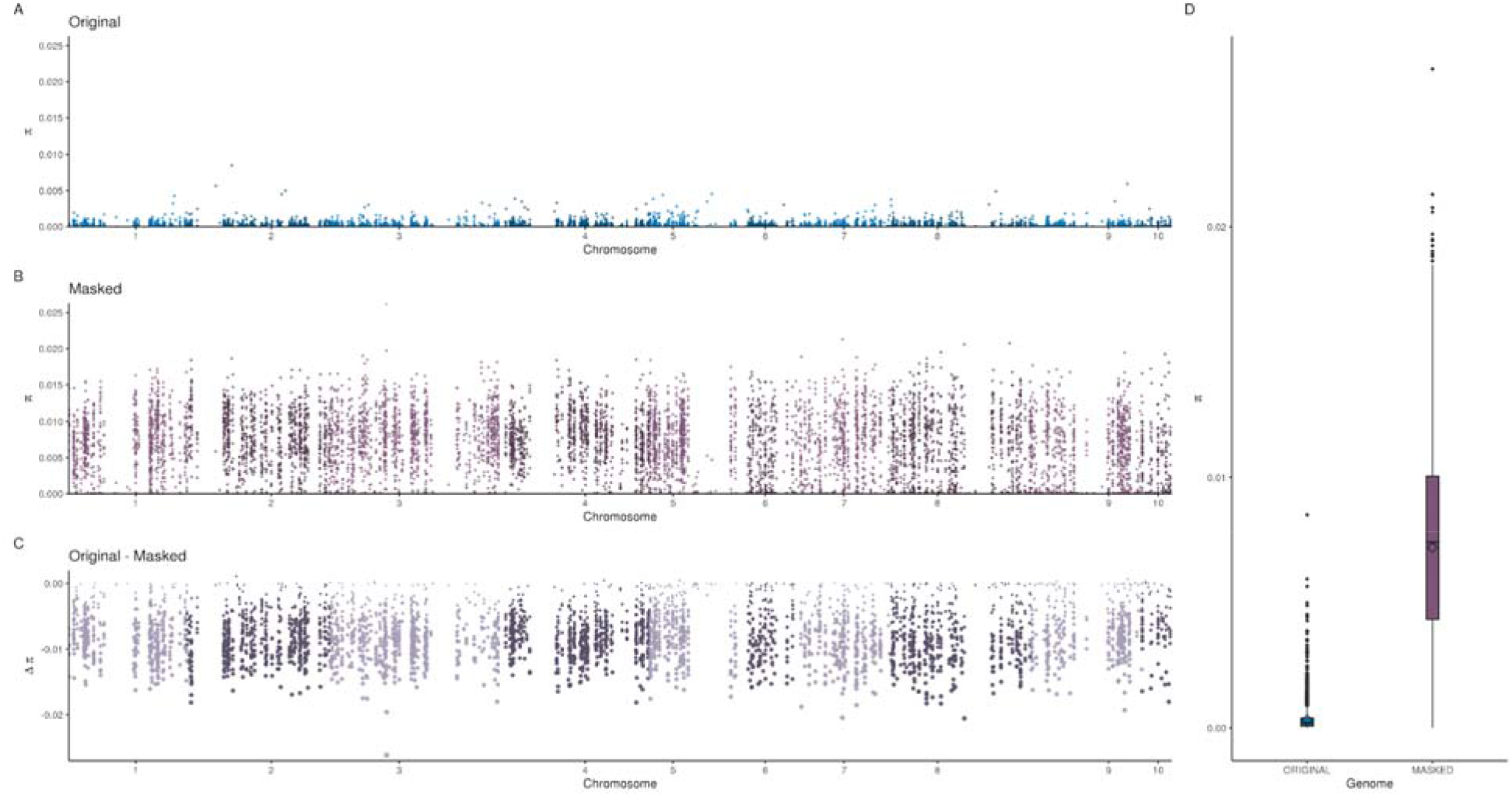
Comparison of nucleotide diversity across the original and haplotig-masked assembly in newly diplotig regions. Panel (A) is values of π averaged across 10kb windows in the original genome that changed to diploid coverage levels after haplotig-masking. Panel (B) panel is values of π averaged across in the same 10kb new diplotig windows across the haplotig-masked genome. Panel (C) is the difference between the original and the haplotig-masked values in 10kb windows across the entire genome with dot size directly related to the distance from zero. Panel (D) is a boxplot of 10kb averaged values between the two genome versions. For all plots, blue is the original genome and purple is the haplotig-masked genome.

### 3.8 Masking haplotigs increases estimates of heterozygosity

The presence of haplotigs in the genome assembly had a more subtle, but statistically significant, effect on measures of heterozygosity compared to nucleotide diversity (Figure 2; Figure S4). When calculated over 10kb windows, the mean heterozygosity of the haplotig-masked genome was 0.140 +/− 0.00249 compared to 0.136 +/− 0.00028 for the original genome assembly. This small difference was statistically significant (*t =* 10.5; df = 74517; *p* < 2.74 x 10^-26^; one-sided t-test). The distribution of heterozygosity differences between genome assemblies showed the greatest difference in new diplotigs (Figure S5). Mean heterozygosity for new diplotigs was significantly higher (*t* = 18.1; df=5980; *p* < 7.36 x 10^-72^; one-sided t-test) in the haplotig-masked assembly (0.142 +/− 0.00046) and had a lower variance compared to the original assembly (0.121 +/− 0.00102; Figure S5).

### 3.9 Haplotig-masking improves the accuracy of estimates of population structure and outlier detection

Overall, the distribution of *F_ST_* values was similar across the two genome versions, with the original assembly having a genome wide average of 0.124 +/− 0.000667 compared to the haplotig-masked genome-wide average of 0.120 +/− 0.000521. This subtle difference was, however, significantly different when evaluated with a one-sided t-test (*t* = -49.2; df=6995427; *p* = 0.000). Estimates of *F_ST_* from the original genome also showed a greater variance than the estimates from the haplotig-masked genome, but in contrast to other population genetic statistics, the distribution of *F_ST_*differences between genome assemblies did not show the greatest difference in new diplotigs (Figure 5; Figure S6). The original genome had a mean *F_ST_* estimate of 0.106 +/− 0.00598 similar to the haplotig-masked genome estimate of 0.109 +/− 9.7 x 10^-5^. Though, this difference was still statistically significant, (one-sided t-test; *t* = 4.81; df=34,218; *p <* 7.71 x 10^-7^).

**Figure 5.**
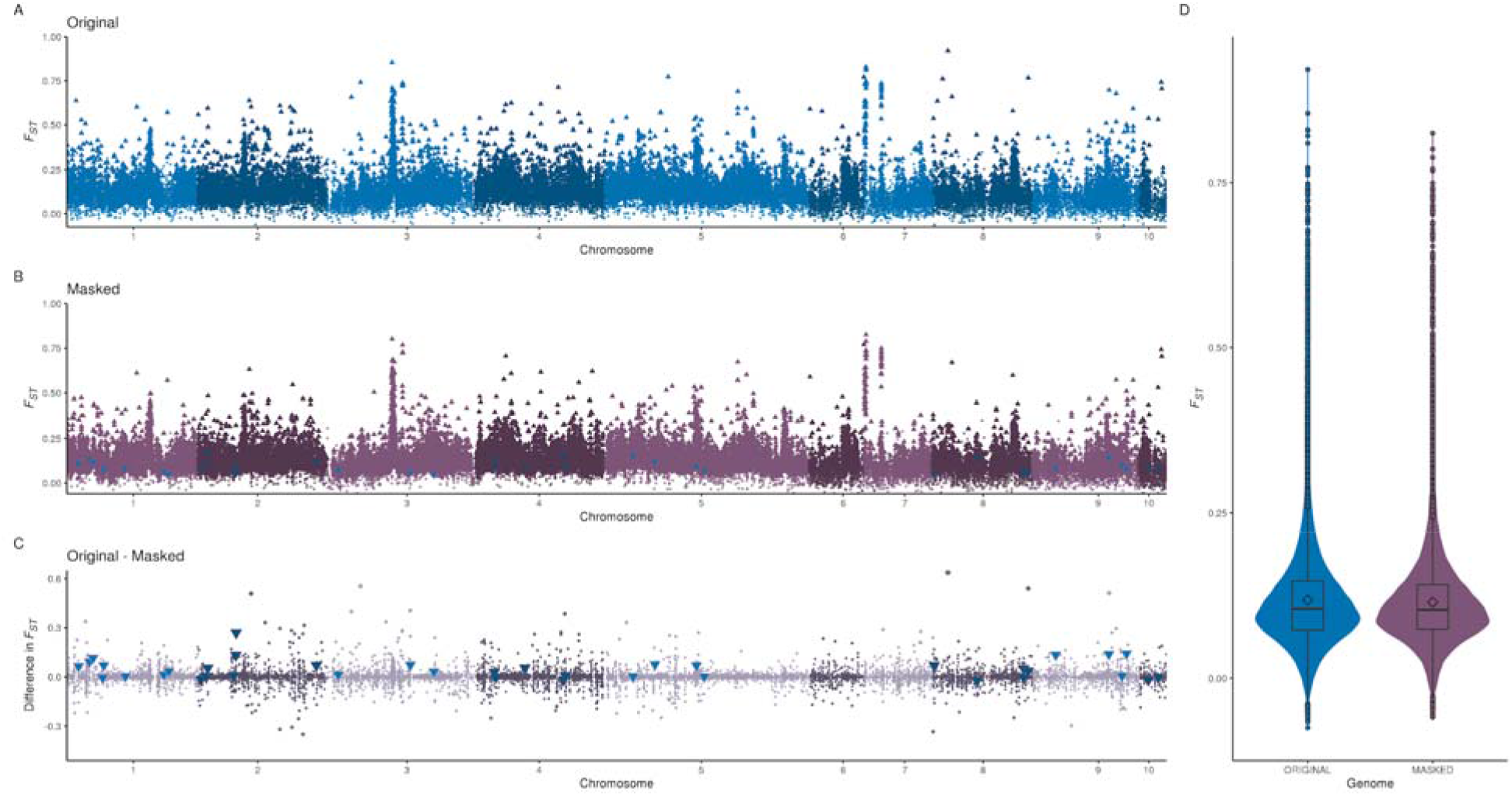
Comparison of estimates of *F_ST_* across the original and haplotig-masked genomes. Panel (A) is values of *F_ST_* averaged across 10kb windows in the original genome. Windows that contained any outlier SNP loci were changed to triangles (16,864 windows). Panel (B) panel is values of *F_ST_* averaged across 10kb windows across the haplotig-masked genome. Windows that contained any outlier SNP loci were changed to triangles (22,182 windows). There were 39 windows with SNPs that were identified as outliers in the original genome analysis but not in the haplotig-masked analysis, and these windows are marked as upside-down blue triangles. Panel (C) is the difference between the original and the haplotig-masked values in 10kb windows across the entire genome with dot size directly related to the distance from zero. There were 39 windows with SNPs that were identified as outliers in the original genome analysis but not in the haplotig-masked analysis, and these windows are marked as upside-down blue triangles. Panel (D) is a violin and boxplot of 10kb averaged values between the two genome versions. For all plots, blue is the original genome and purple is the haplotig-masked genome.

The low structure subset (LSS) showed similar patterns of *F_ST_* values and differences between the two genomes with the original assembly having an average *F_ST_* of 0.0278 +/− 4.8 x 10^-5^ and the haplotig-masked genome having an average *F_ST_* of 0.0257 +/− 3.7 x 10^-5^ (Figure 6). This difference was also significant (one-sided t-test; *t* = -35.6; df = 5980814; *p <* 7.55 x 10^-278^). Again, while variance in estimates was higher in new diplotig regions, the difference in means was less pronounced (Figure S7; original = 0.0181 +/− 0.000427; haplotig-masked = 0.0191+/− 6.78 x 10^-5^) and was not significantly different (one-sided t-test; *t* = 0.779; df =29282; *p* = 0.218).

**Figure 6.**
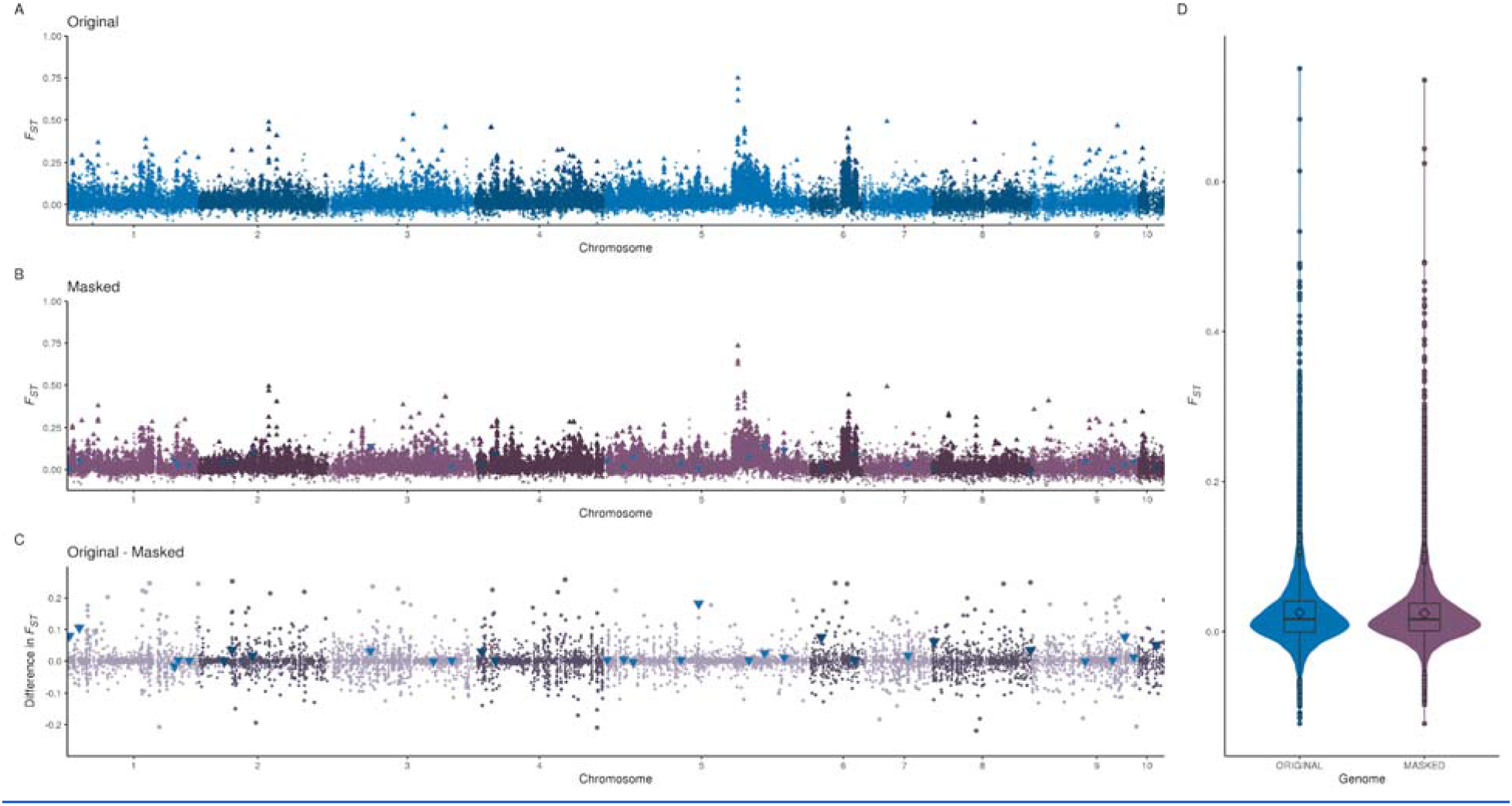
Comparison of estimates of *F_ST_* from the low structure subset (LSS) across the original and haplotig-masked genomes. Estimates of *F_ST_* were calculated using the low structure subset (LSS) to examine how haplotigs affect population structure inference in a lower signal system. Panel (A) is values of *F_ST_* averaged across 10 kb windows in the original genome. Windows that contained any outlier SNP loci were changed to triangles (4,993 windows). Panel (B) panel is values of *F_ST_* averaged across 10kb windows in the haplotig-masked genome. Windows that contained any outlier SNP loci were changed to triangles (3,825 windows). There were 31 windows with SNPs that were identified as outliers in the original genome analysis but not in the haplotig-masked analysis, and these windows are marked as upside-down blue triangles. Panel (C) is the difference between the original and the haplotig-masked values in 10kb windows across the entire genome with dot size directly related to the distance from zero. There were 31 windows with SNPs that were identified as outliers in the original genome analysis but not in the haplotig-masked analysis, and these windows are marked as upside-down blue triangles. Panel (D) is a violin and boxplot of 10kb averaged values between the two genome versions. For all plots, blue is the original genome and purple is the haplotig-masked genome.

Masking haplotigs had a larger effect on outlier loci detection, increasing the number of outliers detected by about 5%. Using a false discovery rate of 5%, OutFLANK detected 158,057 outliers (4.4% of all loci) from data using the original genome assembly contrasted to 257,823 (4.6%) outliers detected from data using the haplotig-masked genome. 2,032 (1.28%) of the outliers detected with the original genome were not called SNPs in the haplotig-masked genome, with 1,249 falling within masked haplotigs. There were an additional 234 (0.15%) outliers from the original genome that were no longer significant in the haplotig-masked genome. Restricting the outlier detection to the LSS, the number of total outliers detected using the original genome was 27,721 (0.96% of all loci) and 38,712 (0.86%) using the haplotig-masked genome. The number of outliers from the original genome that were not present in the masked genome was 260 (0.94%) with 122 (0.44%) loci that were present in the masked haplotigs. 58 (0.21%) outlier loci from the original genome were no longer significant in the masked genome.

### 3.10 Haplotigs reduce the number of detected structural variants

After filtering, the program Delly detected 247,347 different structural variants (SVs) in the original genome compared to 279,390 SVs in the haplotig-masked genome. The haplotig-masked genome had more detections across all categories (Table 4). While the original genome did have less variants detected, the average length was longer for every category of variant (Table 4). When SVs were restricted to only those with no missing data across all individuals, more variants were still detected using the haplotig-masked genome; however, the mean sizes of each variant were either longer for the haplotig-masked genome or nearly identical for with variants detected using the original genome (Supplemental Table 9).

**Table 4.** All Structural Variants Detected.

|  | Type | Number detected | Mean Length | S.E. Length |
| --- | --- | --- | --- | --- |
| Haplotig-Masked | Translocation | 33,012 | NA | NA |
|  | Deletion | 216,912 | 823 | 22 |
|  | Duplication | 15,347 | 14,784 | 464 |
|  | Insertion | 7,310 | 28 | 0 |
|  | Inversion | 6,808 | 118,478 | 2,857 |
| Original | Translocation | 27,838 | NA | NA |
|  | Deletion | 192,011 | 796 | 21 |
|  | Duplication | 13,155 | 17,358 | 585 |
|  | Insertion | 6,351 | 28 | 0 |
|  | Inversion | 7,992 | 243,679 | 3,489 |

### 3.11 Haplotigs may not affect estimates of population frequency of copy number

Delly was also used to call copy number from all samples, and copy number was used to calculate the statistical *V_ST_* across both genomes. Global estimates of *V_ST_*were low for both genome versions (original- 0.0678 +/− 0.00105; haplotig-masked- 0.0663 +/− 0.000916), and they did not differ significantly (two-sided t-test; *t =* -1.10; df = 30,789; *p =* 0.864). Looking at averages across 10kb windows, there was no clear pattern of differentiation between genome versions (Figure S8). The difference was similar when individuals were restricted to the LSS with the original genome version having a global mean estimate of 0.0131 +/− 0.00084 compared to the estimate of 0.0122 +/− 0.00074. This difference was also not statistically significant (two-sided t-test; *t =* -0.83; df = 27,291; *p =* 0.406). Looking across the 10kb windows, the differences between estimates appear to be randomly distributed around zero (Figure S9).

## 4 Discussion

Here, we assembled an annotated, chromosome-level genome for the eastern oyster (*Crassostrea virginica*). The original reference genome, publicly released in 2017, represents one of the most complete and contiguous genomes for a marine invertebrate species. We also present an *ad hoc* method for detecting and masking haplotig sequences in an already published genome that improves coverage, decreases duplicated orthologs while having only nominal impacts on genome completeness. Our results show that masking haplotigs in the eastern oyster genome drastically improved SNP and structural variant discovery. Our results also demonstrate that haplotigs affected population genomic analyses, and that masking haplotigs improved many commonly used population statistics. Taken together, we provide the original assembly and a haplotig-masked genome assembly that will be foundational resources for insights into molluscan adaptation to a changing environment and a valuable resource for the aquaculture industry.

### 4.1 A chromosome-level genome for an important ecosystem engineer and aquaculture and fisheries species

The eastern oyster genome represents a similar level of contiguity and completeness compared to the Pacific oyster (Supplemental table 3; Supplemental Figure S3) and several other published molluscan genomes (Table 2). More broadly, most whole genome assemblies currently available for non-model marine species are fragmented and incomplete (Du et al. 2017; Powell et al. 2018; Gerdol et al. 2020). The *C. gigas* genome was first published in 2012 <u>(Zhang et al., 2012)</u> with updates published by two separate groups in 2021 <u>(Peñaloza et al., 2021; Qi, Li, & Zhang, 2021)</u>. The two updated *C. gigas* assemblies are now chromosome-level along with the eastern oyster genome. The eastern oyster assembly is more complete than the Qi *et al*. (2021) *C. gigas* assembly and is comparable to the Peñaloza *et al*. (2021) *C. gigas* assembly, despite the primary assembly being done several years prior. The haplotig-masked version for *C. virginica* has less duplicates than the Qi *et al*. (2021) *C. gigas* assembly and nearly the same level of duplication as the Peñaloza *et al*. (2021) *C. gigas* assembly, even though haplotigs were masked *post hoc*. The two species have similar numbers of protein coding genes detected, though the higher number for *C. virginica* may have been influenced by the haplotigs present in the original assembly. A shortcoming of the eastern oyster assembly is the mis-assembly of several chromosomal fragments as revealed by the linkage map, which can be corrected in a future assembly. In short, the original eastern oyster genome assembly represents a significant advancement for molluscan and marine invertebrate genomics in its completeness, and the *post hoc* haplotig masking represents a novel way to reduce haplotig sequence without sacrificing genome completeness.

### 4.2 A method for *post hoc* improvement of existing genomic resources

A genome assembly represents an advance in knowledge for any one species as well as a powerful tool for a wide variety of scientific studies. A caveat to a published assembly, or assembly version, is that it represents a single snapshot of a resource that can continually improve over time with both technological improvements and the acquisition of additional high-quality data. Assembly improvements, however, take time, computational and financial resources, and do not always proceed continuously over time or by the same group of researchers. In this study, we have presented a simple methodology for improving existing genome assemblies by masking haplotigs. Looking only at the data generated from the single sequenced genome individual, haplotig-masking greatly improved genome coverage, reducing the number of windows at “haplotig” coverage levels and increasing the windows at “diplotig” coverage levels. Examining read coverage across an exemplar chromosome, areas outside of masked haplotigs that were previously at “haplotig coverage” levels shifted clearly to “diplotig coverage levels”. Most importantly, even though haplotig-masking effectively removed over 100 mb of data, it did not affect genome completeness. The haplotig-masked genome was 99.8% as complete as the original genome but had 85% less duplications (as estimated by BUSCO analysis).

We suspect that the weak point of our method is the breakup of chimeric contigs. The sliding window approach used for this analysis was a simple and successful approach but could be improved by a more sophisticated analysis looking at fine scale patterns of coverage changes statistically, or even incorporating original contig assembly graphs. We also exclusively relied on the program Purge Haplotigs (Roach et al., 2018), and there are new methods and programs, such as HapSolo (Solares et al., 2021) that may be able to offer improvements to our implementation as well.

### 4.3 Haplotigs impact population genomic analyses

An accurate reference genome can enhance our understanding of genome structure, mechanisms promoting genetic diversity and population differentiation, the genetic basis for complex traits, and allow for the investigation of natural and anthropogenic selection (Ekblom & Galindo, 2010; Ellegren, 2014; Fonseca et al., 2016), but mis-assemblies, especially false duplications arising from heterozygosity, can negatively impact SNP discovery (Kelley & Salzberg, 2010; Roach et al., 2018; Solares et al., 2021). We found that the presence of haplotigs in our original assembly greatly impacted SNP discovery, structural variant detection, and significantly impacted all the population genomic statistics that we calculated, including nucleotide diversity, observed heterozygosity, *F_ST_*, and *V_ST_*. The most striking differences were in SNP discovery, where across different data subsets and filtering criteria, the haplotig-masked genome had between 55%-58% more SNPs than the original genome in our resequencing data when any missing data filters were applied, and these differences were most prominent in regions that had coverage increased to diplotig levels after masking. This fits with first-principal expectations that loss of coverage of one allele could lead to a true SNP being mistakenly called an invariant portion of the genome. Interestingly, there was also a small percentage of SNPs (2.28% of all SNPs) that were called in the original genome but not the haplotig-masked. The vast majority were inside haplotigs that were masked, but there were some (20,478; 0.27%) that were in new diplotig regions, indicating that haplotigs do lead to false positive SNPs in one or both allelic copies.

Though discovery was vastly different, SNPs that were genotyped in both genome versions had very high concordance in genotyping (99.62%), though slightly lower in new diplotig regions (98.97%). The high concordance in shared loci lends confidence to any previous results from genomes with potentially small percentages of haplotigs. Though, it should be noted that our results are for genotyped SNPs with moderate (10X-20X) coverage levels per individual and that haplotigs could potentially have a larger effect on low-coverage whole-genome sequencing studies that rely on less coverage per individual and genotype likelihoods instead of genotypes (Lou, Jacobs, Wilder, & Therkildsen, 2021; Lou & Therkildsen, 2022; Matz, 2017).

Perhaps the most important implication of our results is that haplotigs have a large and significant effect on estimates of nucleotide diversity. Estimates of nucleotide diversity were over 50% higher in the haplotig-masked genome, and this is likely directly attributable to the 55% increase in SNP discovery. This is because the more SNPs within a 10kb window, the more likely it is to draw two different haplotypes. The effect of haplotig-masking on nucleotide diversity was most prevalent in new diplotig regions, which in the original genome had estimates of π of close to zero because of a lack of SNPs. For researchers using genomic tools to assess genetic diversity, whether in a conservation application (L. M. Benestan et al., 2016) or a fisheries application (L. Benestan, 2020) of high heterozygosity species, haplotig-masking of existing genomic resources should be a critical step before population-level assessment.

In contrast to nucleotide diversity, our results suggest that haplotigs have only a minor impact on overall estimates of heterozygosity. The haplotig-masked genome did have significantly higher estimates of heterozygosity than the original genome, but the difference was approximately 3% across the whole genome and 17.5% in new diplotig regions. The subtle difference is likely due to the high concordance of shared genotypes between the two genome versions, as observed heterozygosity is simply a proportion of variable loci that are heterozygous and not greatly affected by differences in SNP discovery.

The results from our population structure and outlier detection analyses were more nuanced. Estimates of global *F_ST_* were 3.33% larger, on average, estimated from the original genome version compared to the haplotig-masked genome using the full data set; however, there was a much larger difference in estimates when using the lower structure subset with the original genome having estimates that were 8.2% larger on average than the haplotig-masked genome. Not only did haplotigs inflate estimates of *F_ST_*, but they also increased the variance of those estimates; the original genome estimates had a standard error 28% higher than the haplotig-masked genome in the full dataset and 29% in the LSS. The increase in variance had implications for outlier detection. 1,483 out of 158,057 outliers that were significant in the full dataset analysis using the original genome were either not significant or no longer present in the analysis with the haplotig-masked genome. For the lower structure subset, there were 180 outliers out of 27,721 that were missing or non-significant. The differences we observed were small but consistent, and there are some potential caveats to our analysis. First, our per locality sample sizes were small (only six individuals per population), and this likely affected the power we had to detect small allele frequency differences. Second, our SNPs were genotyped at moderate coverage levels and highly filtered, tolerating no missing data. Missing data may potentially interact with haplotig effects to alter allele frequencies, but we did not test this in our study. Lastly, we only examined global population structure and pairwise estimates may have different patterns due to smaller sample sizes and potentially even smaller background levels of population structure. Taken all together, haplotigs have a small but consistent and significant effect on estimates of population structure, and there are still potential haplotig effects that remain unknown.

Understanding the role of structural variants in adaptation and population structure has taken a more prominent role in molecular ecology (Bazzicalupo et al., 2020; Mérot, Oomen, Tigano, & Wellenreuther, 2020; Nelson et al., 2019; Prunier et al., 2019; Wellenreuther, Mérot, Berdan, & Bernatchez, 2019). Our analysis indicates that while haplotigs do affect structural variant detection and discovery, haplotigs do not affect estimates of differentiation based on copy number variation. We found that analysis with the haplotig-masked genome found more structural variants than the original genome, though all variants were smaller on average in the haplotig-masked genome. The differences in length did virtually disappear if the analysis was restricted to only variant calls without missing data. Estimates of *V_ST_* were virtually identical between genome versions for the full data set, and there was a small but non-significant difference when using the lower structure subset. We only used one program to estimate both copy number variation and identify structural variants, and our analysis may have benefited from a stand-alone estimation of copy number variation. We also only had moderate levels of per sample coverage (10X-20X) this may have limited our power to detect differences in copy number variation between genome versions.

## 5 Conclusion

In this manuscript, we present a chromosome-level genome assembly of the eastern oyster (*Crassostrea virginica*), and we describe an ad-hoc method for masking haplotig sequences, including chimeric contigs, within an existing assembly. We show that haplotig masking improves read mapping, genome coverage, and SNP discovery. The haplotig-masked genome greatly reduced duplicated orthologs, while still maintaining one of the highest levels of genome completeness and continuity for molluscan genomes. Resequencing data shows that haplotig-masking greatly improves estimates of nucleotide diversity and offers subtle but significant improvements to estimates of heterozygosity population structure, and outlier detection. The eastern oyster genome (original and haplotig-masked) will help support both fundamental, applied, and conservation research on a critical ecosystem species and one of the largest aquaculture species in North America.

## Acknowledgements

We would like to thank K. Markey Lundgren from the USDA ARS lab for technical support and Gregory DeBrosse for oyster caring. We also thank members of the Eastern Oyster Genome Consortium and the Eastern Oyster Breeding Consortium for intellectual input during data analysis. This work was funded by USDA NIFA AFRI Award#2015-67016-22942 to MGC and WW, a USDA NRSP8 aquaculture award to DP and MGC, the College of Environmental and Life Sciences at the University of Rhode Island, and USDA-NIFA Hatch Program projects 1017848/RI0019-H020 and 1021665/NJ30401. This is contribution 001 from the Puritz Lab of Marine Evolutionary Ecology.

## Data Accessibility

Raw sequencing reads for the original assembly and haplotig masking are archived at the NCBI Short Read Archive (BioProject: PRJNA376014). Individual Accession numbers are also provided in Supplemental Table S2. Raw demultiplexed sequencing reads for the population resequencing analysis are archived at the NCBI Short Read Archive (BioProject: PRJNA951786). Additionally, all data, fully reproducible code for the haplotig masking and genome version comparisons, and the haplotig-masked genome can be found at the “Haplotig-masked Eastern Oyster Genome” repository (http://doi.org/10.17605/OSF.IO/9SM76). The masked genome is archived through Zenodo available at <u>DOI: 10.5281/zenodo.7799622</u>.

## Author Contributions

For the original genome assembly and resequencing project, X.G., M.H., D.P., W.W., and M.G-C. designed the project and contributed intellectually, X.G., M.H., and M.L. collected samples, Y.H., L.H, S.J., M.L., P.M., T.M, D.P., E.W., and C.T provided technical help including DNA and RNA isolation and performing various other experiments. X.G., M.H., Y.H., L.H., P.M., T.M., D.P., E.S.R., C.T., W.W, E.W., and M.G-C. contributed to the genome assembly, annotation, and curation. J.B.P developed the haplotig-masking protocol and performed all analyses comparing the two genome versions. J.B.P, X.G., M.H., K.L., D.P. H.Z., and M.G-C contributed intellectually to the haplotig-masked genome comparison and wrote the manuscript. J.B.P, X.G., D.P., and M.G-C. provided funding for this work.

## References

1. Albertin, C. B., Simakov, O., Mitros, T., Wang, Z. Y., Pungor, J. R., Edsinger-Gonzales, E.,… Rokhsar, D. S. (2015). The octopus genome and the evolution of cephalopod neural and morphological novelties. Nature, 524(7564), 220–224.

2. Allen, S. K., Jr, Small, J. M., & Kube, P. D. (2021). Genetic parameters for Crassostrea virginica and their application to family-based breeding in the mid-Atlantic, USA. *Aquaculture (Amsterdam*, Netherlands), 538(736578), 736578.

3. Bazzicalupo, A. L., Ruytinx, J., Ke, Y.-H., Coninx, L., Colpaert, J. V., Nguyen, N. H.,… Branco, S. (2020). Fungal heavy metal adaptation through single nucleotide polymorphisms and copy-number variation. Molecular Ecology. doi: 10.1111/mec.15618

4. Benestan, L. (2020). Population Genomics Applied to Fishery Management and Conservation. In M. F. Oleksiak & O. P. Rajora (Eds.), Population Genomics: Marine Organisms (pp. 399– 421). Cham: Springer International Publishing.

5. Benestan, L. M., Ferchaud, A.-L., Hohenlohe, P. A., Garner, B. A., Naylor, G. J. P., Baums, I. B.,… Luikart, G. (2016). Conservation genomics of natural and managed populations: building a conceptual and practical framework. Molecular Ecology, 25(13), 2967–2977.

6. Bhagwat, M., Young, L., & Robison, R. R. (2012). Using BLAT to find sequence similarity in closely related genomes. Current Protocols in Bioinformatics / Editoral Board, Andreas D. Baxevanis… [et Al.], Chapter 10, 10.8.1-10.8.24.

7. Bickhart, D. M., Rosen, B. D., Koren, S., Sayre, B. L., Hastie, A. R., Chan, S.,… Smith, T. P. L. (2017). Single-molecule sequencing and chromatin conformation capture enable de novo reference assembly of the domestic goat genome. Nature Genetics, 49(4), 643–650.

8. Boetzer, M., & Pirovano, W. (2014). SSPACE-LongRead: scaffolding bacterial draft genomes using long read sequence information. BMC Bioinformatics, 15, 211.

9. Botwright, N. A., Zhao, M., Wang, T., McWilliam, S., Colgrave, M. L., Hlinka, O.,… Cummins, S. F. (2019). Greenlip Abalone (Haliotis laevigata) Genome and Protein Analysis Provides Insights into Maturation and Spawning. G3, 9(10), 3067–3078.

10. Buroker, N. E. (1983). Population genetics of the American oyster Crassostrea virginica along the Atlantic coast and the Gulf of Mexico. Marine Biology, 75(1), 99–112.

11. Catchen, J., Amores, A., & Bassham, S. (2020). Chromonomer: A Tool Set for Repairing and Enhancing Assembled Genomes Through Integration of Genetic Maps and Conserved Synteny. G3, 10(11), 4115–4128.

12. Chen, N. (2004). Using RepeatMasker to identify repetitive elements in genomic sequences. Current Protocols in Bioinformatics / Editoral Board, Andreas D. Baxevanis… [et Al.], Chapter 4(1), Unit 4.10.

13. Chen, S., Zhou, Y., Chen, Y., & Gu, J. (2018). fastp: an ultra-fast all-in-one FASTQ preprocessor. Bioinformatics, 34(17), i884–i890.

14. Chin, C.-S., Alexander, D. H., Marks, P., Klammer, A. A., Drake, J., Heiner, C.,… Korlach, J. (2013). Nonhybrid, finished microbial genome assemblies from long-read SMRT sequencing data. Nature Methods, 10(6), 563–569.

15. Danecek, P., Auton, A., Abecasis, G., Albers, C. A., Banks, E., DePristo, M. A.,… Durbin, R. (2011). The variant call format and VCFtools. Bioinformatics, 27(15), 2156–2158.

16. Danecek, P., Bonfield, J. K., Liddle, J., Marshall, J., Ohan, V., Pollard, M. O.,… Li, H. (2021). Twelve years of SAMtools and BCFtools. GigaScience, 10(2). doi: 10.1093/gigascience/giab008

17. Du, X., Fan, G., Jiao, Y., Zhang, H., Guo, X., Huang, R.,… Liu, X. (2017). The pearl oyster Pinctada fucata martensii genome and multi-omic analyses provide insights into biomineralization. GigaScience, 6(8), 1–12.

18. Ekblom, R., & Galindo, J. (2010). Applications of next generation sequencing in molecular ecology of non-model organisms. Heredity, 107(1), 1–15.

19. Ellegren, H. (2014). Genome sequencing and population genomics in non-model organisms. Trends in Ecology & Evolution, 29(1), 51–63.

20. Fonseca, R., Albrechtsen, A., Sibbesen, J. A., Maretty, L., Zepeda-mendoza, M. L., Campos, P. F., & Pereira, R. J. (2016). Next-generation biology: Sequencing and data analysis approaches for non-model organisms. Marine Genomics. doi: 10.1016/j.margen.2016.04.012

21. Garrison, E. (2016). Vcflib, a simple C++ library for parsing and manipulating VCF files.

22. Garrison, E. and Marth, G. (2012). Haplotype-Based Variant Detection from Short-Read Sequencing.

23. Ghanayim, A. (2013). Iterative Referencing for Improving the Interpretation of Dna Sequence Data.

22. Garrison, Erik, & Marth, G. (2012). Haplotype-based variant detection from short-read sequencing -- Free bayes -- Variant Calling -- Longranger. ArXiv Preprint ArXiv:1207.3907. doi: arXiv:1207.3907 [q-bio.GN]

23. Gerdol, M., Moreira, R., Cruz, F., Gómez-Garrido, J., Vlasova, A., Rosani, U.,… Figueras, A. (2020). Massive gene presence-absence variation shapes an open pan-genome in the Mediterranean mussel. Genome Biology, 21(1), 275.

24. Gómez-Chiarri, M., Warren, W. C., Guo, X., & Proestou, D. (2015). Developing tools for the study of molluscan immunity: The sequencing of the genome of the eastern oyster, Crassostrea virginica. Fish & Shellfish Immunology, 46(1), 2–4.

25. Grabherr, M. G., Haas, B. J., Yassour, M., Levin, J. Z., Thompson, D. A., Amit, I.,… Regev, A. (2011). Full-length transcriptome assembly from RNA-Seq data without a reference genome. Nature Biotechnology, 29(7), 644–652.

26. Grabowski, J., Conrad, F., & James, J. (2012). Economic Valuation of Ecosystem Services Provided by Oyster Reefs. Bioscience, 62(10), 900–909.

27. Guo, X. (2021). Genetics in shellfish culture. In S. Shumway (Ed.), Molluscan Shellfish Aquaculture: A Practical Guide. 5m Books Ltd.

28. Guo, X., Hershberger, W. K., Cooper, K., & Chew, K. K. (1993). Artificial gynogenesis with ultraviolet light-irradiated sperm in the Pacific oyster, Crassostrea gigas. I. Induction and survival. Aquaculture, 113(3), 201–214.

29. He, Y. (2012). Identification and application of disease-resistance markers in the eastern oyster (Crassostrea virginica) (Ph. D.). Ocean University of China.

30. Hoover, Cindi A., and Patrick M. Gaffney. (2005). Geographic variation in nuclear genes of the eastern oyster, Crassostrea virginica Gmelin. J. Shellfish Res., 24(1), 103–112.

31. Institute, B. (2016). Picard tools. Broad Institute, GitHub repository.

32. James Kent, W. (2002). BLAT—The BLAST-Like Alignment Tool. Genome Research, 12(4), 656–664.

33. Kajitani, R., Toshimoto, K., Noguchi, H., Toyoda, A., Ogura, Y., Okuno, M.,… Itoh, T. (2014). Efficient de novo assembly of highly heterozygous genomes from whole-genome shotgun short reads. Genome Research, 24(8), 1384–1395.

34. Karl, S. A., & Avise, J. C. (1992). Balancing selection at allozyme loci in oysters: implications from nuclear RFLPs. Science, 256(5053), 100–102.

35. Kelley, D. R., & Salzberg, S. L. (2010). Detection and correction of false segmental duplications caused by genome mis-assembly. Genome Biology, 11(3), R28.

36. Kim, B.-M., Kang, S., Ahn, D.-H., Jung, S.-H., Rhee, H., Yoo, J. S.,… An, H. S. (2018). The genome of common long-arm octopus Octopus minor. GigaScience, 7(11). doi: 10.1093/gigascience/giy119

37. Li, H. (2018). Minimap2: pairwise alignment for nucleotide sequences. Bioinformatics, 34(18), 3094–3100.

38. Li, H. (2021). New strategies to improve minimap2 alignment accuracy. Bioinformatics. doi:10.1093/bioinformatics/btab705

39. Li, H., & Durbin, R. (2010). Fast and accurate long-read alignment with Burrows-Wheeler transform. Bioinformatics, 26(5), 589–595.

40. Li, H., Handsaker, B., Wysoker, A., Fennell, T., Ruan, J., Homer, N.,… Durbin, R. (2009). The Sequence Alignment/Map format and SAMtools. Bioinformatics, 25(16), 2078–2079.

41. Lou, R. N., Jacobs, A., Wilder, A. P., & Therkildsen, N. O. (2021). A beginner’s guide to low-coverage whole genome sequencing for population genomics. Molecular Ecology, 30(23), 5966–5993.

42. Lou, R. N., & Therkildsen, N. O. (2022). Batch effects in population genomic studies with low-coverage whole genome sequencing data: Causes, detection and mitigation. Molecular Ecology Resources, 22(5), 1678–1692.

43. Masonbrink, R. E., Purcell, C. M., Boles, S. E., Whitehead, A., Hyde, J. R., Seetharam, A. S., & Severin, A. J. (2019). An Annotated Genome for Haliotis rufescens (Red Abalone) and Resequenced Green, Pink, Pinto, Black, and White Abalone Species. Genome Biology and Evolution, 11(2), 431–438.

44. Matz, M. V. (2017). Fantastic Beasts and How To Sequence Them: Ecological Genomics for Obscure Model Organisms. Trends in Genetics: TIG, xx, 1–12.

45. Mérot, C., Oomen, R. A., Tigano, A., & Wellenreuther, M. (2020). A Roadmap for Understanding the Evolutionary Significance of Structural Genomic Variation. Trends in Ecology & Evolution, 35(7), 561–572.

46. Murgarella, M., Puiu, D., Novoa, B., Figueras, A., Posada, D., & Canchaya, C. (2016). Correction: A First Insight into the Genome of the Filter-Feeder Mussel Mytilus galloprovincialis. PloS One, 11(7), e0160081.

47. Nelson, T. C., Monnahan, P. J., McIntosh, M. K., Anderson, K., MacArthur-Waltz, E., Finseth, F. R.,… Fishman, L. (2019). Extreme copy number variation at a tRNA ligase gene affecting phenology and fitness in yellow monkeyflowers. Molecular Ecology, 28(6), 1460–1475.

48. Peñaloza, C., Gutierrez, A. P., Eöry, L., Wang, S., Guo, X., Archibald, A. L.,… Houston, R. D. (2021). A chromosome-level genome assembly for the Pacific oyster Crassostrea gigas. GigaScience, 10(3). doi: 10.1093/gigascience/giab020

49. Powell, D., Subramanian, S., Suwansa-Ard, S., Zhao, M., O’Connor, W., Raftos, D., & Elizur, A. (2018). The genome of the oyster Saccostrea offers insight into the environmental resilience of bivalves. DNA Research: An International Journal for Rapid Publication of Reports on Genes and Genomes, 25(6), 655–665.

50. Privé, F., Aschard, H., Ziyatdinov, A., & Blum, M. G. B. (2018). Efficient analysis of large-scale genome-wide data with two R packages: bigstatsr and bigsnpr. Bioinformatics, 34(16), 2781–2787.

51. Pruitt, K. D., Tatusova, T., Brown, G. R., & Maglott, D. R. (2012). NCBI Reference Sequences (RefSeq): current status, new features and genome annotation policy. Nucleic Acids Research, 40(Database issue), D130–5.

52. Prunier, J., Giguère, I., Ryan, N., Guy, R., Soolanayakanahally, R., Isabel, N.,… Porth, I. (2019). Gene copy number variations involved in balsam poplar (Populus balsamifera L.) adaptive variations. Molecular Ecology, 28(6), 1476–1490.

53. Pryszcz, L. P., & Gabaldón, T. (2016). Redundans: an assembly pipeline for highly heterozygous genomes. Nucleic Acids Research, 44(12), e113.

54. Puritz, J. B., Hollenbeck, C. M., & Gold, J. R. (2014). dDocent: a RADseq, variant-calling pipeline designed for population genomics of non-model organisms. PeerJ, 2, e431.

55. Puritz, J. B., Zhao, H., Guo, X., Hare, M. P., He, Y., LaPeyre, J.,… Gomez-Chiarri, M. (2022). Nucleotide and structural polymorphisms of the eastern oyster genome paint a mosaic of divergence, selection, and human impacts. BioRxiv.

56. Qi, H., Li, L., & Zhang, G. (2021). Construction of a chromosome-level genome and variation map for the Pacific oyster Crassostrea gigas. Molecular Ecology Resources, 21(5), 1670–1685.

57. Quinlan, A. R., & Hall, I. M. (2010). BEDTools: a flexible suite of utilities for comparing genomic features. Bioinformatics, 26(6), 841–842.

58. R Development Core Team. (2008). R: A Language and Environment for Statistical Computing. Vienna, Austria: R Foundation for Statistical Computing. Retrieved from http://www.r-project.org

59. Rausch, T., Zichner, T., Schlattl, A., Stütz, A. M., Benes, V., & Korbel, J. O. (2012). DELLY: structural variant discovery by integrated paired-end and split-read analysis. Bioinformatics, 28(18), i333–i339.

60. Redon, R., Ishikawa, S., Fitch, K. R., Feuk, L., Perry, G. H., Andrews, T. D.,… Hurles, M. E. (2006). Global variation in copy number in the human genome. Nature, 444(7118), 444– 454.

61. Reeb, C. A., & Avise, J. C. (1990). A genetic discontinuity in a continuously distributed species: mitochondrial DNA in the American oyster, Crassostrea virginica. Genetics, 124(2), 397– 406.

62. Roach, M. J., Schmidt, S. A., & Borneman, A. R. (2018). Purge Haplotigs: Allelic contig reassignment for third-gen diploid genome assemblies. BMC Bioinformatics, 19(1), 1–10.

63. Safonova, Y., Bankevich, A., & Pevzner, P. A. (2015). dipSPAdes: Assembler for Highly Polymorphic Diploid Genomes. Journal of Computational Biology: A Journal of Computational Molecular Cell Biology, 22(6), 528–545.

64. Schwartz, S., James Kent, W., Smit, A., Zhang, Z., Baertsch, R., Hardison, R. C.,… Miller, W. (2003). Human–Mouse Alignments with BLASTZ. Genome Research, 13(1), 103–107.

65. Seppey, M., Manni, M., & Zdobnov, E. M. (2019). BUSCO: Assessing Genome Assembly and Annotation Completeness. Methods in Molecular Biology, 1962, 227–245.

66. Simakov, O., Marletaz, F., Cho, S.-J., Edsinger-Gonzales, E., Havlak, P., Hellsten, U.,… Rokhsar, D. S. (2013). Insights into bilaterian evolution from three spiralian genomes. Nature, 493(7433), 526–531.

67. Simão, F. A., Waterhouse, R. M., Ioannidis, P., Kriventseva, E. V., & Zdobnov, E. M. (2015). BUSCO: assessing genome assembly and annotation completeness with single-copy orthologs. Bioinformatics, 31(19), 3210–3212.

68. Solares, E. A., Tao, Y., Long, A. D., & Gaut, B. S. (2021). HapSolo: an optimization approach for removing secondary haplotigs during diploid genome assembly and scaffolding. BMC Bioinformatics, 22(1), 9.

69. Steenwyk, J. L., Soghigian, J. S., Perfect, J. R., & Gibbons, J. G. (2016). Copy number variation contributes to cryptic genetic variation in outbreak lineages of Cryptococcus gattii from the North American Pacific Northwest. BMC Genomics, 17, 700.

70. Thompson, P. A., Guo, M.-X., & Harrison, P. J. (1996). Nutritional value of diets that vary in fatty acid composition for larval Pacific oysters (Crassostrea gigas). Aquaculture, 143(3), 379–391.

71. van Dijk, E. L., Auger, H., Jaszczyszyn, Y., & Thermes, C. (2014). Ten years of next-generation sequencing technology. Trends in Genetics: TIG, 30(9), 418–426.

72. Varney, R. L., Galindo-Sánchez, C. E., Cruz, P., & Gaffney, P. M. (2009). Population Genetics of the Eastern Oyster Crassostrea virginica (Gmelin, 1791) in the Gulf of Mexico. Journal of Shellfish Research, 28(4), 855–864.

73. Vinson, J. P., Jaffe, D. B., O’Neill, K., Karlsson, E. K., Stange-Thomann, N., Anderson, S.,… Lander, E. S. (2005). Assembly of polymorphic genomes: algorithms and application to Ciona savignyi. Genome Research, 15(8), 1127–1135.

74. Walker, B. J., Abeel, T., Shea, T., Priest, M., Abouelliel, A., Sakthikumar, S.,… Earl, A. M. (2014). Pilon: an integrated tool for comprehensive microbial variant detection and genome assembly improvement. PloS One, 9(11), e112963.

75. Wang, Y., Wang, X., Wang, A., & Guo, X. (2010). A 16-microsatellite multiplex assay for parentage assignment in the eastern oyster (Crassostrea virginica Gmelin). Aquaculture, 308, S28–S33.

76. Wellenreuther, M., Mérot, C., Berdan, E., & Bernatchez, L. (2019). Going beyond SNPs: the role of structural genomic variants in adaptive evolution and species diversification. Molecular Ecology, (February), 1203–1209.

77. Whitlock, M. C., & Lotterhos, K. E. (2015). Reliable Detection of Loci Responsible for Local Adaptation: Inference of a Null Model through Trimming the Distribution of *F* _ST_. The American Naturalist, 186(S1), S24–S36.

78. Wickham, H. (2016). Data Analysis. In H. Wickham (Ed.), ggplot2: Elegant Graphics for Data Analysis (pp. 189–201). Cham: Springer International Publishing.

79. Yang, J.-L., Feng, D.-D., Liu, J., Xu, J.-K., Chen, K., Li, Y.-F.,… Lu, Y. (2021). Chromosome-level genome assembly of the hard-shelled mussel Mytilus coruscus, a widely distributed species from the temperate areas of East Asia. GigaScience, 10(4). doi: 10.1093/gigascience/giab024

80. Zarrella, I., Herten, K., Maes, G. E., Tai, S., Yang, M., Seuntjens, E.,… Fiorito, G. (2019). The survey and reference assisted assembly of the Octopus vulgaris genome. Scientific Data, 6(1), 13.

81. Zhang, G., Fang, X., Guo, X., Li, L., Luo, R., Xu, F.,… Wang, J. (2012). The oyster genome reveals stress adaptation and complexity of shell formation. Nature, 490(7418), 49–54.

