## Supplemental Information for "A second unveiling: haplotig masking of the eastern oyster genome improves population-level inference" (from another version of this preprint: version 2, 2022-12-17)

### Supplemental Table 1. Evaluation of heterozygosity in gynogenetic progeny using microsatellite markers.

**pl**

| Group | Aged of oysters at test | #Microsat. Loci | Parent | Progeny (n) |
| --- | --- | --- | --- | --- |
| XGHG1 | 12 months | 11 | 0.182 | 0.042 (24) |
|  | 22 months | 15 | 0.333 | 0.187 (9) |
|  | Average |  | 0.258 | 0.115 (33) |
| Wild oysters | 12 months | 11 | - | 0.642 (60) |

### Supplemental Table 2. C. virginica sequence sources for de novo assembly analyses.

| **Table S2.** C. virginica sequence sources for *de novo* assembly analyses. | | |
| --- | --- | --- |
| Genetic resources | NCBI BioProject ID | NCBI Accession Number |
| Illumina WGS | PRJNA376014 | SRX2652900 |
| SMRT sequences | PRJNA376014 | SRX3142107; SRX3142106; SRX3142105 |
| Assembly | PRJNA376014 | GCA_002022765.4 |
| Mitochondria genome |  |  |
| Transcriptome resources |  |  |
| adductor muscle |  | SRX2652901 |
| digestive |  | SRX2652897 |
| gill |  | SRX2652892 |
| mantle |  | SRX2652893 |

### Supplemental Table 3. Genus Crassostrea genome assembly comparisons.

|  |  | **Total size (Mb)** | **N50 contig length (Mb)** | **N50 scaffold length (Mb)** |  |
| --- | --- | --- | --- | --- | --- |
| **Genus species** | **Assembly version** |  |  |  | **Total contigs** |
| *C. virginica* | C virginica 3.0 | 684 | 1.97 | 75 | 669 |
| *C. gigas* | cgigas uk roslin v1 | 647 | 1.56 | 58 | 783 |
| *C. gigas* | ASM1103280v1 | 586 | 3.12 | 61 | 301 |
| a. All assembly metrics are derived from the NCBI assembly databases. | | | | |  |
| b. N50 contig or scaffold length is defined as the length N for which 50% of all bases in the assembly sequences L are in a sequence of length N >L. | | | | | |

### Supplemental Table 4. Representative gene annotation metrics for oyster genomes of genus Crassostrea

| **Supplemental Table 4.** Representative gene annotation metrics for oyster genomes of genus Crassostrea | | | | | |
| --- | --- | --- | --- | --- | --- |
|  |  |  | Protein-coding genes | Total ncRNA | mRNAs |
| Common name | Species | Assembled version |  |  |  |
| Eastern oyster | *C. virginica* | C virginica 3.0 | 34,596 | 4,320 | 60,201 |
| Pacific oyster | *C. gigas* | cgigas uk roslin v1 | 31,371 | 6,837 | 63,341 |

### Supplemental Table 5. Summary of the repeat element classification.

| **Supplementary Table 5.** Summary of the repeat element classification. | |
| --- | --- |
| Repeat category | % Genome-wide |
| Repeat masked | 32.86 |
| Tandem repeats | 7.74 |
| SINE | 1.65 |
| LINE | 1.75 |
| LTR elements | 0.77 |
| DNA elements | 5.42 |
| Unclassified | 22.38 |
| Small RNA | 0.23 |
| Simple repeats | 0.28 |
| Low complexity | 0.5 |
| Total Masking | 40.6 |
| Windowmasker | 36 |

### Supplemental Table 6. Correspondence between assembled chromosomes (Chrs) and linkage groups (LGs) of the genetic map.

| **Chr** | **Reference** | **Total length** | **Correspondence between LGs and assembled chromosomes** |
| --- | --- | --- | --- |
| 1 | [NC_035780.1](https://www.ncbi.nlm.nih.gov/nuccore/NC_035780.1) | 65,668,440 | LG1 = Chr1 |
| 2 | [NC_035781.1](https://www.ncbi.nlm.nih.gov/nuccore/NC_035781.1) | 61,752,955 | LG2 = Chr2 |
| 3 | [NC_035782.1](https://www.ncbi.nlm.nih.gov/nuccore/NC_035782.1) | 77,061,148 | LG3 = Chr3 |
| 4 | [NC_035783.1](https://www.ncbi.nlm.nih.gov/nuccore/NC_035783.1) | 59,691,872 | LG4 = Chr4 |
| 5 | [NC_035784.1](https://www.ncbi.nlm.nih.gov/nuccore/NC_035784.1) | 98,698,416 | LG5 = Chr5:3,019,001-61,274,182; Chr5:61,754,194-97,730,408 = LG6 |
| 6 | [NC_035785.1](https://www.ncbi.nlm.nih.gov/nuccore/NC_035785.1) | 51,258,098 | LG6 = Chr6:30,846,534-44,031,109 + Chr5:61,754,194-97,730,408; Chr6:207,314-29,816,260 + Chr6:44,251,733-51,022,757 = LG7 |
| 7 | [NC_035786.1](https://www.ncbi.nlm.nih.gov/nuccore/NC_035786.1) | 57,830,854 | LG7 = Chr7 + Chr6:207,314-29,816,260 + Chr6:44,251,733-51,022,757 |
| 8 | [NC_035787.1](https://www.ncbi.nlm.nih.gov/nuccore/NC_035787.1) | 75,944,018 | LG8 = Chr8 |
| 9 | [NC_035788.1](https://www.ncbi.nlm.nih.gov/nuccore/NC_035788.1) | 104,168,038 | LG9 = Chr9:1,481,457-5,573,926 + Chr9:48,528,867-90,680,673 |
| 10 | [NC_035789.1](https://www.ncbi.nlm.nih.gov/nuccore/NC_035789.1) | 32,650,045 | LG10 = Chr10 + Chr9:7,020,053-47,804,347 + Chr9:91,590,937-103,471,324 |
| *The genetic map was constructed with 4006 RAD-seq markers segregating in a F2 family with 115 progeny (He, 2012). | | | |

### Supplemental Table 8. Sequencing and mapping information for both genome versions

| **Samples** | | | **Haplotig-Masked Genome** | | | **Original Genome** | | |
| --- | --- | --- | --- | --- | --- | --- | --- | --- |
| **Sample** | **Raw Reads** | **Trimmed Reads** | **Total Mappings** | **Duplicates Detected** | **Post-filtering Mappings** | **Total Mappings** | **Duplicates Detected** | **Post-filtering Mappings** |
| CL_1 | 61756328 | 60515962 | 58638967 | 3678309 | 42779510 | 58792994 | 3610775 | 41417721 |
| CL_2 | 99774458 | 97882558 | 94839920 | 5785436 | 69565701 | 95094136 | 5674605 | 67388718 |
| CL_3 | 96409828 | 92943592 | 89955536 | 5728720 | 64336032 | 90181900 | 5599577 | 62143042 |
| CL_4 | 57385744 | 56122644 | 54376130 | 3384483 | 39568166 | 54522577 | 3320852 | 38244358 |
| CL_5 | 51825176 | 49808752 | 48188040 | 2849220 | 34477409 | 48314973 | 2791733 | 33320632 |
| CL_6 | 66656598 | 64735606 | 62427500 | 3773383 | 45608980 | 62609727 | 3702975 | 44146068 |
| CLP_1 | 59304898 | 57214842 | 55227242 | 3427383 | 40531929 | 55388084 | 3371633 | 39433861 |
| CLP_2 | 70623926 | 68392668 | 65942791 | 4459706 | 48158612 | 66130606 | 4388533 | 46883686 |
| CLP_3 | 98389272 | 94723928 | 91658037 | 5644819 | 65954366 | 91898433 | 5527982 | 64118169 |
| CLP_4 | 69924272 | 67476732 | 65189185 | 4573182 | 46891439 | 65370779 | 4476255 | 45558283 |
| CLP_5 | 84034792 | 81291734 | 78410746 | 5500463 | 57109934 | 78650383 | 5411942 | 55597354 |
| CLP_6 | 97689796 | 93990226 | 90836203 | 6891883 | 65385413 | 91102894 | 6766237 | 63595134 |
| CS_1 | 76082270 | 73729270 | 71169316 | 4784194 | 51922862 | 71383322 | 4699062 | 50494450 |
| CS_2 | 72128202 | 69666346 | 67261377 | 4260850 | 49558329 | 67474877 | 4188203 | 48226389 |
| CS_3 | 68535568 | 66784598 | 64869270 | 4143160 | 48776174 | 65047586 | 4075430 | 47583908 |
| CS_5 | 75917232 | 74059434 | 71760922 | 4403643 | 52503142 | 71952425 | 4315009 | 51008408 |
| CS_6 | 64004962 | 61921988 | 59831080 | 4224183 | 43734808 | 60006329 | 4158286 | 42557317 |
| CS_7 | 69777108 | 68030738 | 65877071 | 4122388 | 48391958 | 66063487 | 4053809 | 47037945 |
| DEBY_1 | 70340066 | 68129214 | 65984679 | 4204859 | 48013858 | 66164945 | 4122462 | 46680477 |
| DEBY_2 | 61820396 | 59393626 | 57501832 | 3569039 | 41033127 | 57645855 | 3488757 | 39723953 |
| DEBY_3 | 90050284 | 88202564 | 85646456 | 5163796 | 62737184 | 85852948 | 5060844 | 60888813 |
| DEBY_4 | 54919768 | 52712608 | 51030035 | 3250463 | 37069517 | 51166462 | 3190492 | 35998396 |
| DEBY_5 | 66912230 | 65379052 | 63242334 | 4157130 | 46365584 | 63419266 | 4087871 | 45066905 |
| DEBY_6 | 69395392 | 67069200 | 64901712 | 4409876 | 46839037 | 65076101 | 4327613 | 45517708 |
| HC_1 | 77293636 | 74906388 | 72425035 | 5835437 | 52186665 | 72629897 | 5739699 | 50785966 |
| HC_3 | 63112478 | 61557868 | 59524843 | 4346896 | 43322163 | 59697110 | 4280145 | 42155592 |
| HC_4 | 79754444 | 77660564 | 75074963 | 5501949 | 54611969 | 75281513 | 5417016 | 53141136 |
| HC_5 | 90033632 | 87108604 | 84111972 | 5616048 | 61600005 | 84359499 | 5513394 | 59972504 |
| HC_6 | 56048260 | 54559620 | 52342661 | 3436210 | 38373096 | 52494916 | 3382841 | 37334935 |
| HC_7 | 154599222 | 148626848 | 143848308 | 8804482 | 103414438 | 144224018 | 8543578 | 100379268 |
| HC_VA_1 | 85940448 | 82636266 | 79780756 | 5891673 | 57681262 | 80020832 | 5782645 | 56110025 |
| HC_VA_2 | 82902150 | 79755572 | 77204226 | 5527680 | 55580071 | 77424184 | 5419862 | 54056327 |
| HC_VA_3 | 63890928 | 61749506 | 59643898 | 3294694 | 44391895 | 59825571 | 3234312 | 43190499 |
| HC_VA_4 | 69606580 | 67810232 | 65615532 | 3866507 | 48549698 | 65798257 | 3800871 | 47198225 |
| HC_VA_5 | 79755810 | 77137600 | 74474190 | 5370365 | 54248567 | 74697466 | 5278626 | 52802055 |
| HC_VA_6 | 85495276 | 82438178 | 79705400 | 6118503 | 57718329 | 79931281 | 6015385 | 56165305 |
| HG_HG0F2 | 81488634 | 79609440 | 78557006 | 4643260 | 65052524 | 78736248 | 4611315 | 64650437 |
| HG_HG2F1 | 107490278 | 104562398 | 102891704 | 7466672 | 78838999 | 103124360 | 7327313 | 78504587 |
| HG_HG2M5 | 65635204 | 64848234 | 64299271 | 4276150 | 49711444 | 64374580 | 4177329 | 48847629 |
| HI_1 | 78587340 | 76634458 | 74051860 | 5107038 | 54685883 | 74261901 | 5028043 | 53294696 |
| HI_2 | 118354864 | 114591096 | 110920625 | 8030373 | 81152249 | 111234084 | 7884228 | 79161418 |
| HI_3 | 84232622 | 81870440 | 79113148 | 5748565 | 57429647 | 79351366 | 5646539 | 55889409 |
| HI_4 | 77417784 | 75521270 | 73141247 | 5281685 | 53400559 | 73355963 | 5197788 | 51967585 |
| HI_5 | 104038488 | 101912426 | 98927966 | 6886611 | 73404346 | 99199192 | 6778858 | 71632775 |
| HI_6 | 78733988 | 76799260 | 74276699 | 5147995 | 54206331 | 74487694 | 5059072 | 52708938 |
| LM_1_pool | 66969314 | 64216034 | 61769102 | 4561219 | 41620194 | 61945674 | 4447766 | 40046145 |
| LM_3 | 92486240 | 88551324 | 84623488 | 5074628 | 58897350 | 84914650 | 4959058 | 56800294 |
| LM_4 | 172376322 | 166674276 | 159956314 | 9420105 | 111641861 | 160434045 | 9156388 | 107615488 |
| LM_7 | 75118986 | 72981782 | 70198561 | 4065303 | 50087656 | 70417228 | 3981776 | 48400022 |
| LM_8 | 66373270 | 64272486 | 61604541 | 3484785 | 43154467 | 61812572 | 3411318 | 41622951 |
| LOLA_1 | 92988582 | 90136906 | 86935582 | 5759948 | 63012286 | 87165449 | 5649931 | 61104961 |
| LOLA_2 | 67427816 | 65354868 | 63047433 | 3849653 | 46353891 | 63224125 | 3782013 | 44966092 |
| LOLA_3 | 70995646 | 68916992 | 66510965 | 4070465 | 48922303 | 66700322 | 4000451 | 47445730 |
| LOLA_4 | 81746434 | 78906376 | 76174709 | 5454965 | 54236799 | 76384280 | 5344988 | 52618845 |
| LOLA_5 | 68704786 | 66973404 | 64653570 | 4445880 | 46785631 | 64829436 | 4366570 | 45353384 |
| LOLA_6 | 77903354 | 75647386 | 73198687 | 4979400 | 52565163 | 73387480 | 4885298 | 50951248 |
| NEH_1 | 62806886 | 61154804 | 59252416 | 3971608 | 44112015 | 59405816 | 3909288 | 42992110 |
| NEH_2 | 73087636 | 71245062 | 69207906 | 4771209 | 51989894 | 69393229 | 4700616 | 50783839 |
| NEH_3 | 103930046 | 101703694 | 99125055 | 7295972 | 73002332 | 99361278 | 7168900 | 71127295 |
| NEH_4 | 116701620 | 112592462 | 109362296 | 7223266 | 80135399 | 109629772 | 7087686 | 78065309 |
| NEH_5 | 64394368 | 62707068 | 61129630 | 4349320 | 45033834 | 61269552 | 4272742 | 43835463 |
| NEH_6 | 74547538 | 72648536 | 70714452 | 5006221 | 51309059 | 70882423 | 4915119 | 49913115 |
| NG_NH0H4 | 70988650 | 69279808 | 67299078 | 4214975 | 50693549 | 67484057 | 4151741 | 49481592 |
| NG_NH2F6 | 64297714 | 62713746 | 61018545 | 4418426 | 45139472 | 61180405 | 4350881 | 44029108 |
| NG_NH2F8 | 67198670 | 65281042 | 63378230 | 4452766 | 46583461 | 63556844 | 4382631 | 45463335 |
| NG_NH2M1 | 79668514 | 77463988 | 75181995 | 5423725 | 55065837 | 75388350 | 5333799 | 53744777 |
| OBOYS2_1 | 65043442 | 63682580 | 61624990 | 3846448 | 45074106 | 61793637 | 3778954 | 43686408 |
| OBOYS2_2 | 70055162 | 68520216 | 66194378 | 4446615 | 48035537 | 66384164 | 4368918 | 46560160 |
| OBOYS2_3 | 82925952 | 79746152 | 77181900 | 4533366 | 55447440 | 77382928 | 4433726 | 53631719 |
| OBOYS2_4 | 56605592 | 54813194 | 52683361 | 3429467 | 37926799 | 52830134 | 3367466 | 36716936 |
| OBOYS2_5 | 71635564 | 70124370 | 67818633 | 4475680 | 49208853 | 68006676 | 4395533 | 47682026 |
| OBOYS2_6 | 107126892 | 104962500 | 101646882 | 6572431 | 74053208 | 101917756 | 6444063 | 71747597 |
| SL_1 | 89147578 | 86978844 | 83523180 | 5065708 | 61078007 | 83758253 | 4964751 | 59236119 |
| SL_2 | 59238048 | 57637436 | 55584184 | 3426593 | 40568507 | 55743266 | 3367617 | 39321066 |
| SL_3 | 93694418 | 91917534 | 88796974 | 5843833 | 64699574 | 89046054 | 5736281 | 62756520 |
| SL_4 | 61696302 | 60440650 | 58460518 | 3522935 | 42836220 | 58619664 | 3458060 | 41497645 |
| SL_5 | 99844072 | 97535802 | 94365829 | 6174554 | 68891226 | 94635842 | 6059350 | 66839731 |
| SL_6 | 76386656 | 74247366 | 71756216 | 4338350 | 52284751 | 71954115 | 4250919 | 50686062 |
| SM_10 | 83663278 | 81453140 | 78799232 | 5721628 | 57257531 | 79013818 | 5625996 | 55694728 |
| SM_11 | 63381898 | 61514580 | 57785915 | 3795790 | 41969957 | 57961692 | 3731820 | 40784674 |
| SM_12 | 71593066 | 69750482 | 67385513 | 4770414 | 49074758 | 67582251 | 4694585 | 47754405 |
| SM_7 | 95076544 | 93039658 | 90035576 | 7358208 | 64816528 | 90280377 | 7242642 | 63082190 |
| SM_8 | 71562510 | 69681052 | 67407309 | 4892705 | 49016663 | 67597002 | 4804618 | 47633629 |
| SM_9 | 71256028 | 69432742 | 67144886 | 4394338 | 49081725 | 67335706 | 4318734 | 47704245 |
| UMFS_1 | 64143576 | 62429272 | 60777720 | 4095873 | 43909923 | 60922190 | 4007567 | 42646132 |
| UMFS_2 | 76203502 | 74877078 | 72822380 | 5144437 | 53057441 | 73014343 | 5046314 | 51562047 |
| UMFS_3 | 99560248 | 97380516 | 94283095 | 6361244 | 69210032 | 94546796 | 6248119 | 67368457 |
| UMFS_4 | 94236090 | 91852914 | 88970604 | 6190590 | 64826142 | 89222947 | 6070836 | 62995616 |
| UMFS_5 | 71144408 | 68855926 | 66650390 | 4162606 | 48791995 | 66834913 | 4074048 | 47415877 |
| UMFS_6 | 72412090 | 70347494 | 67922828 | 4481380 | 49830314 | 68131172 | 4404356 | 48508718 |

### Supplemental Table 9. All Structural Variants Detected with no missing data allowed

|  | Type | Number detected | Mean Length | S.E. Length |
| --- | --- | --- | --- | --- |
| Haplotig-Masked | Translocation | 1153 | N/A | N/A |
|  | Deletion | 21158 | 354 | 38 |
|  | Duplication | 1611 | 5996 | 997 |
|  | Insertion | 413 | 29 | 0 |
|  | Inversion | 519 | 52839 | 6819 |
| Original | Translocation | 749 | N/A | N/A |
|  | Deletion | 16953 | 355 | 42 |
|  | Duplication | 1269 | 5544 | 1094 |
|  | Insertion | 336 | 29 | 0 |
|  | Inversion | 428 | 57553 | 8023 |

### Figure S1- Comparison of coverage across original and haplotig masked assembly

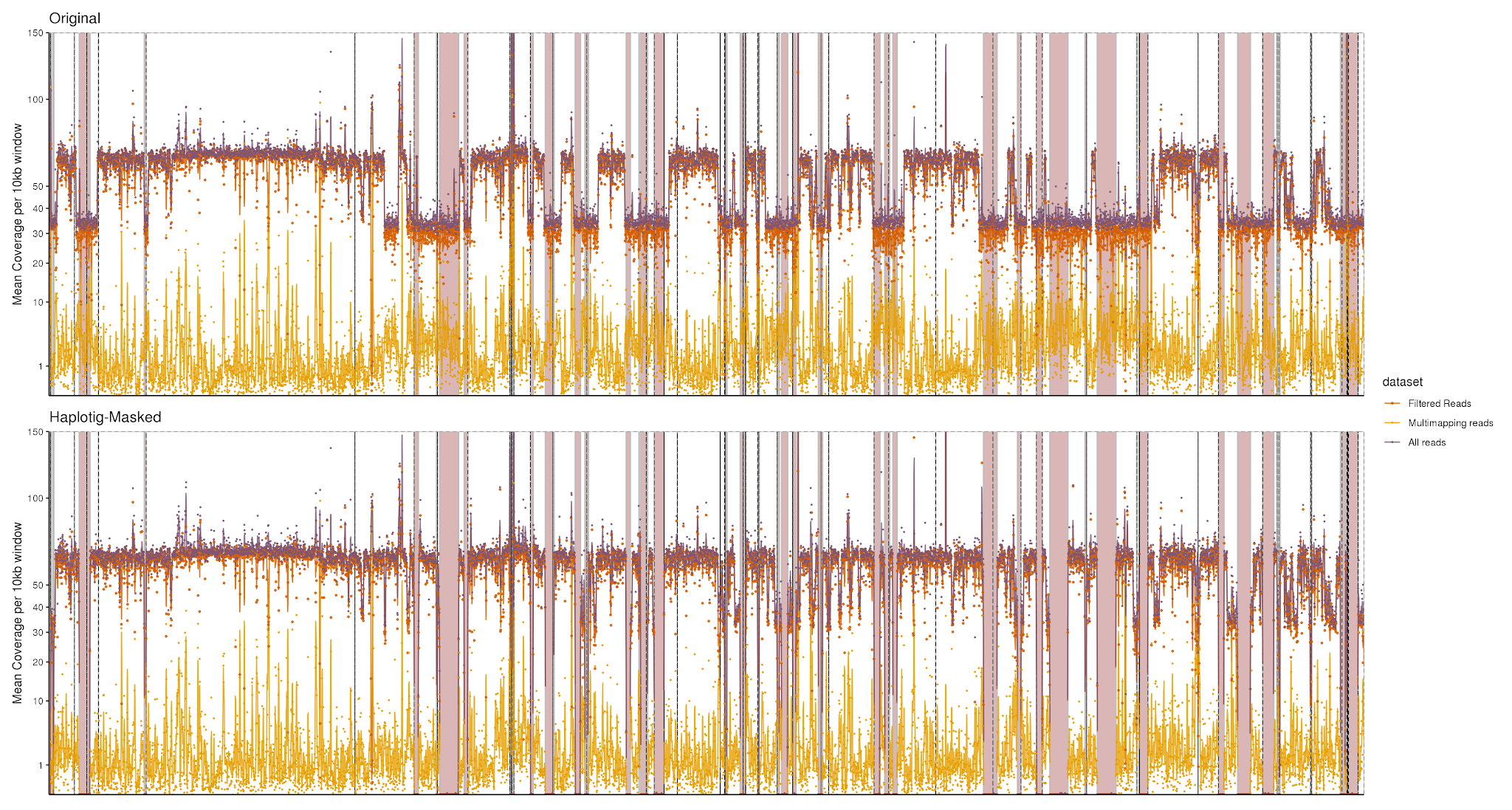
<https://github.com/The-Eastern-Oyster-Genome-Project/2022_Eastern_Oyster_Haplotig_Masked_Genome/raw/main/Comparative_Analysis/Output/Figures/Supplemental/FigureS1.CoverageChromosome2.png>

**Figure S1- Comparison of coverage across original and haplotig-masked assembly.** Across both assemblies, coverage of reads from the original genome individual was averaged across ten kilobase windows of Chromosome 2 (NC_03578.1). The top panel is the original genome and the lower panel is the haplotig-masked genome. Points are the values per 10 kb window with lines drawn as rolling three point averages The color of each line and point differentiates the filtering applied to reads before coverage calculations. Purple is total mappings excluding duplicates and primary alignments. Orange is coverage of reads post-filtration for duplicates, low-quality mapping, multi-mapping reads, and secondary alignments. Yellow are reads that mapped with equal probability to multiple locations in the genome. Areas along the chromosome shaded in dark red were identified as haplotigs and therefore have no data for the haplotig-masked genome.

### Figure S2- Busco results from both genome versions

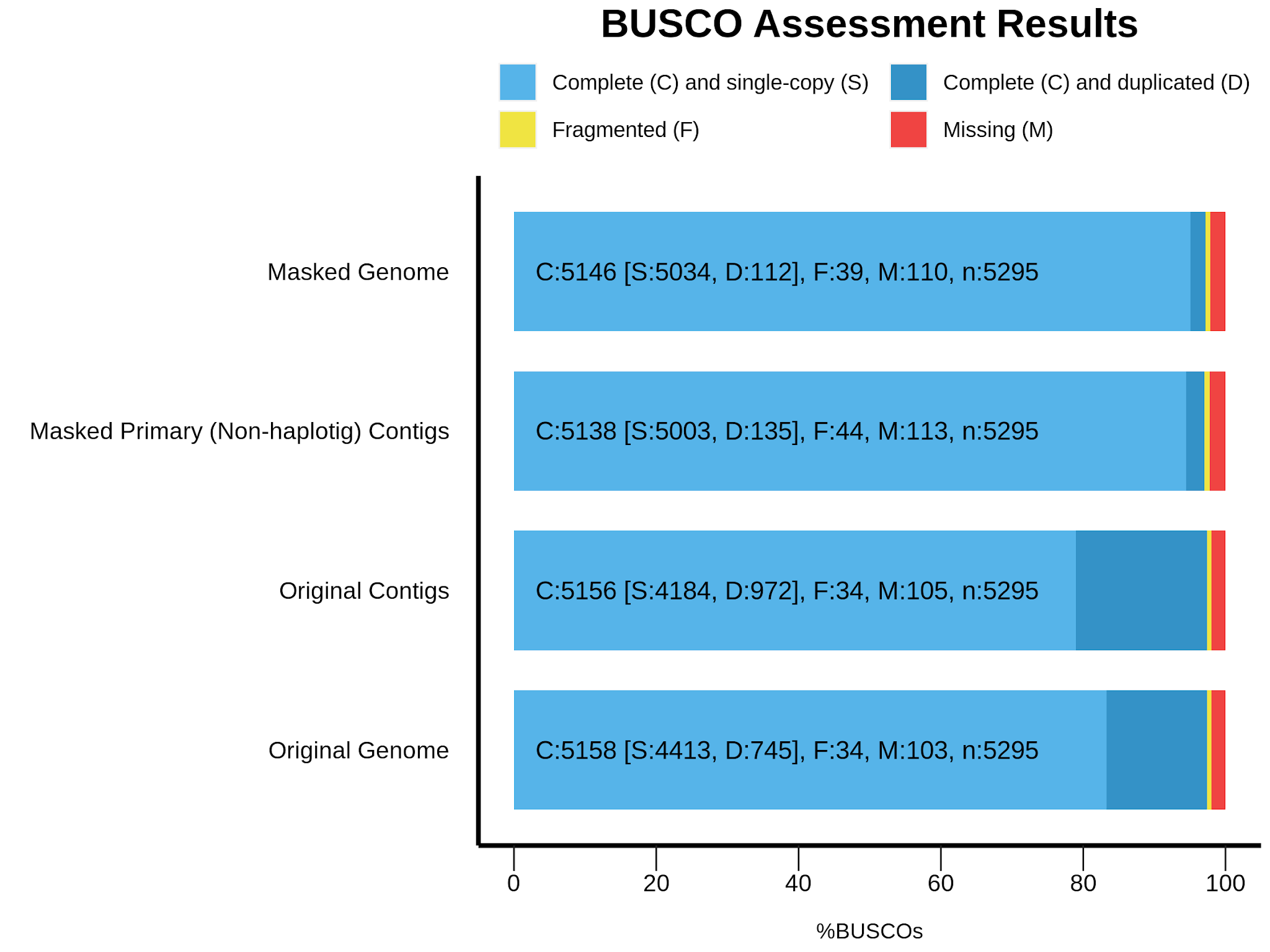
<https://github.com/The-Eastern-Oyster-Genome-Project/2022_Eastern_Oyster_Haplotig_Masked_Genome/raw/main/Comparative_Analysis/Output/BUSCO/Summaries/busco_figure.png>

**Figure S2- BUSCO results plotted as a bar graph.** The results of the four different BUSCO analyses using the Mollusca specific ortholog database. Complete and single-copy orthologs are in light blue, complete and duplicated are in dark blue, fragment orthologs are in yellow, and missing orthologs are in red. All assemblies had over 97% completeness, and haplotip masking dropped duplication rates down from 14.06% to 2.11%.

### Figure S3- Busco results in comparison to *C. gigas*

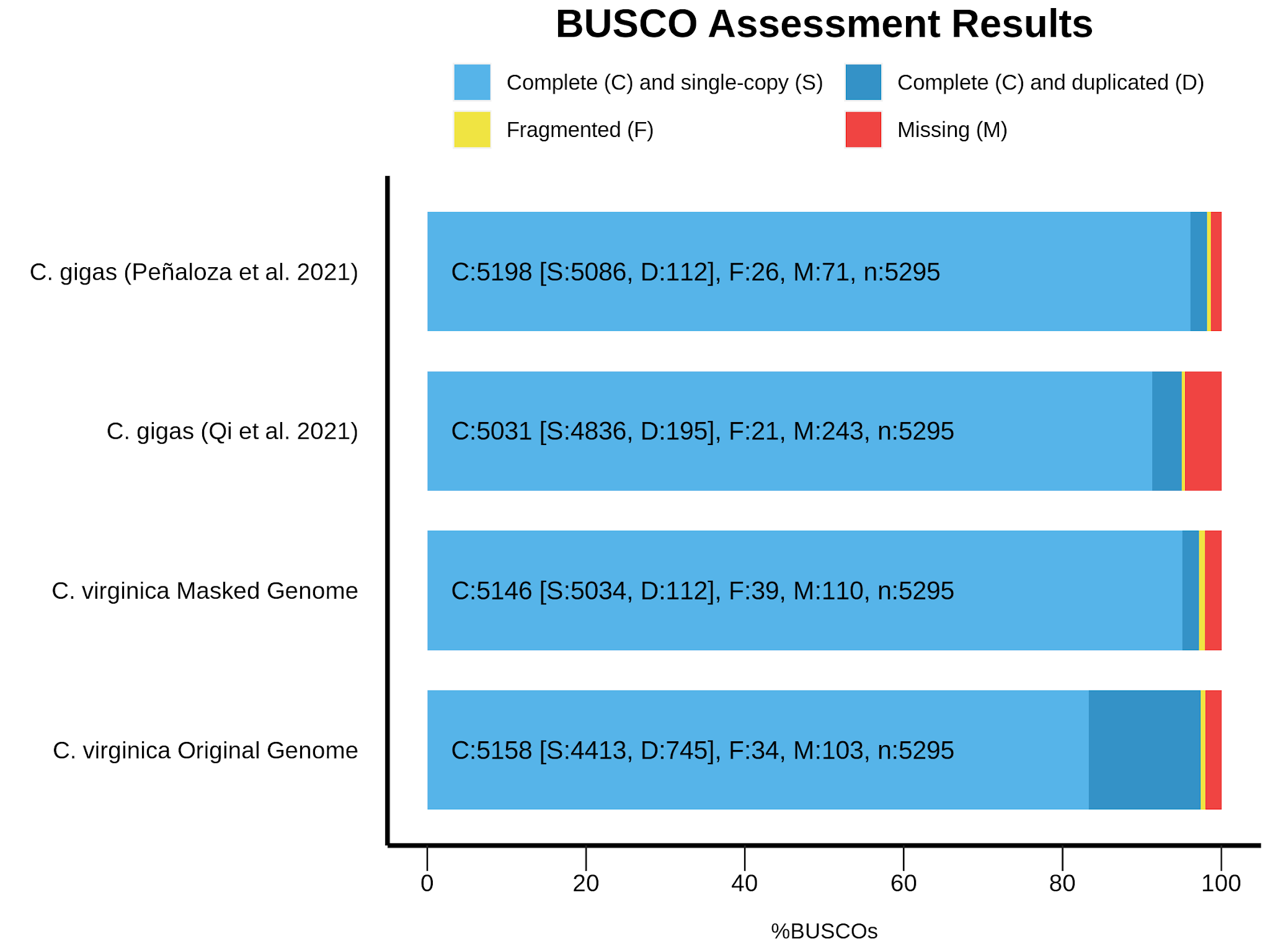
<https://github.com/The-Eastern-Oyster-Genome-Project/2022_Eastern_Oyster_Haplotig_Masked_Genome/raw/main/Comparative_Analysis/Output/BUSCO/Summaries/busco_figure.png>**Figure S3- Busco results in comparison to *C. gigas*.**The results of BUSCO analyses using the Mollusca specific ortholog database comparing the two *C. virginica* assemblies with *C. gigas*. Complete and single-copy orthologs are in light blue, complete and duplicated are in dark blue, fragment orthologs are in yellow, and missing orthologs are in red.

### Figure S4- The effect of haplotigs in estimates of heterozygosity across the genome

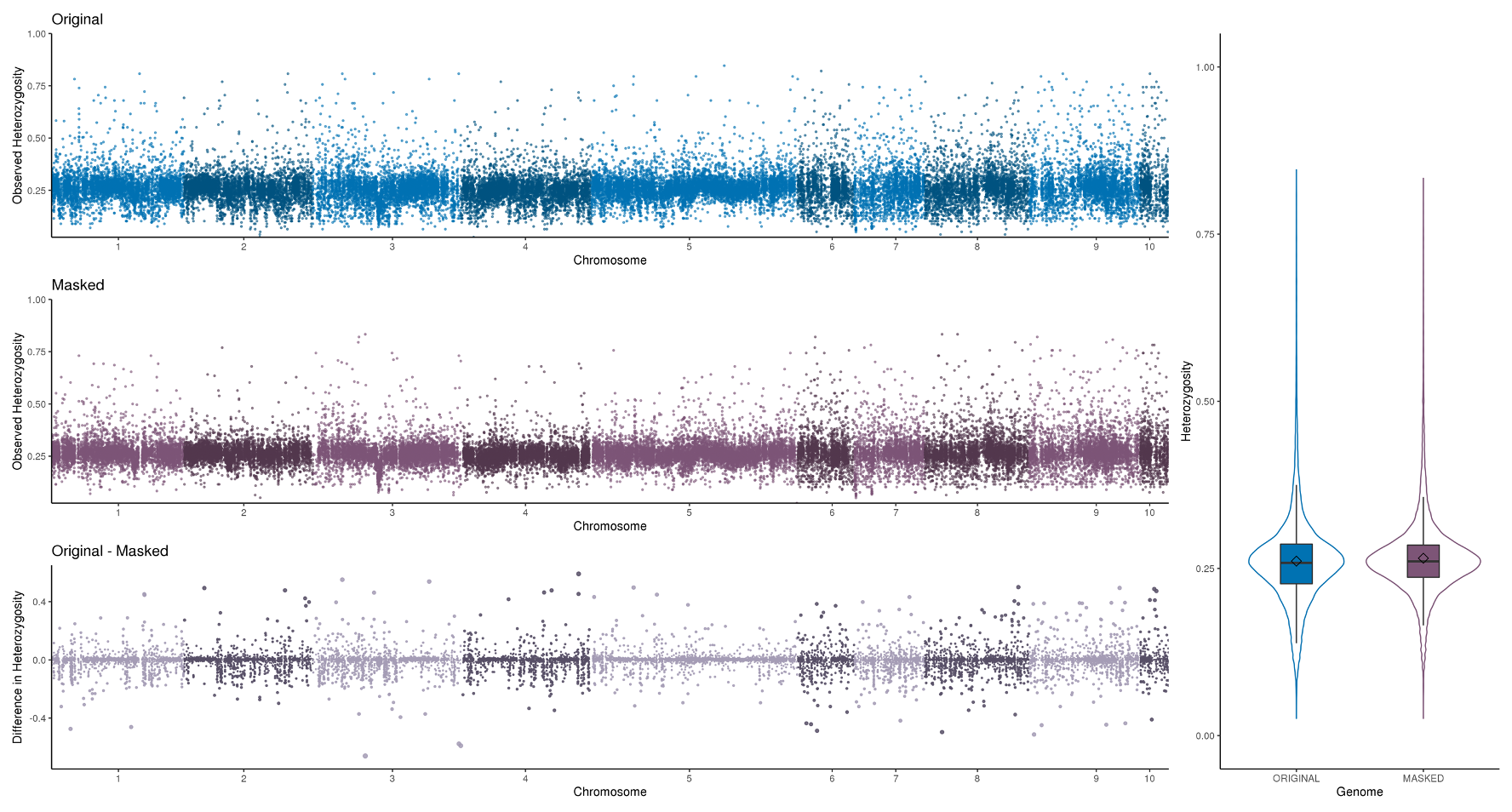

**Figure S4- Comparison of heterozygosity across the original and haplotig-masked assembly.** Panel (A) is values of *H*_O_ averaged across 10 kb windows across the original genome. Panel (B) panel is values of *H*_O_ averaged across 10 kb windows across the haplotig-masked genome. Panel (C) is the difference between the original and the haplotig-masked values in 10 kb windows across the entire genome. Panel (D) is a violin and boxplot of 10kb averaged values between the two genome versions. For all plots, blue is the original genome and purple is the haplotig-masked genome.

### Figure S5- The effect of haplotigs in estimates of heterozygosity across diplotigs

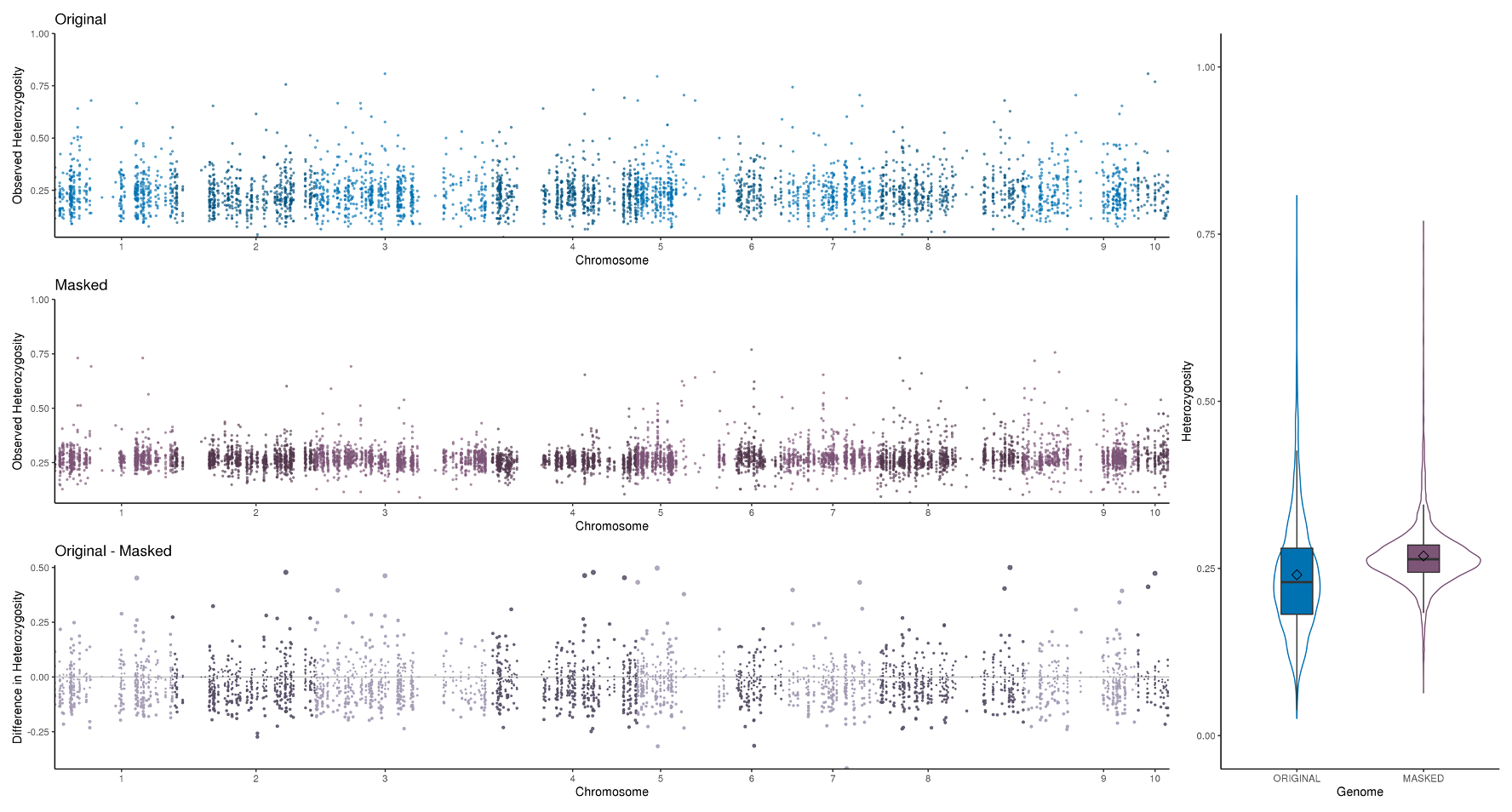

**Figure S5- Comparison of heterozygosity across the original and haplotig-masked assembly in newly diplotigous regions.** Panel (A) is values of *H*_O_ averaged across 10 kb windows in the original genome that changed to diplotigous coverage levels after haplotig-masking. Panel (B) panel is values of *H*_O_ averaged across in the same 10 kb newly diplogtigous windows across the haplotig-masked genome. Panel (C) is the difference between the original and the haplotig-masked values in 10 kb windows across the entire genome with dot size directly related to the distance from zero. Panel (D) is a boxplot of 10kb averaged values between the two genome versions. For all plots, blue is the original genome and purple is the haplotig-masked genome

### Figure S6- Comparison of estimates of FST across new diplotigous regions in the original and haplotig-masked genomes

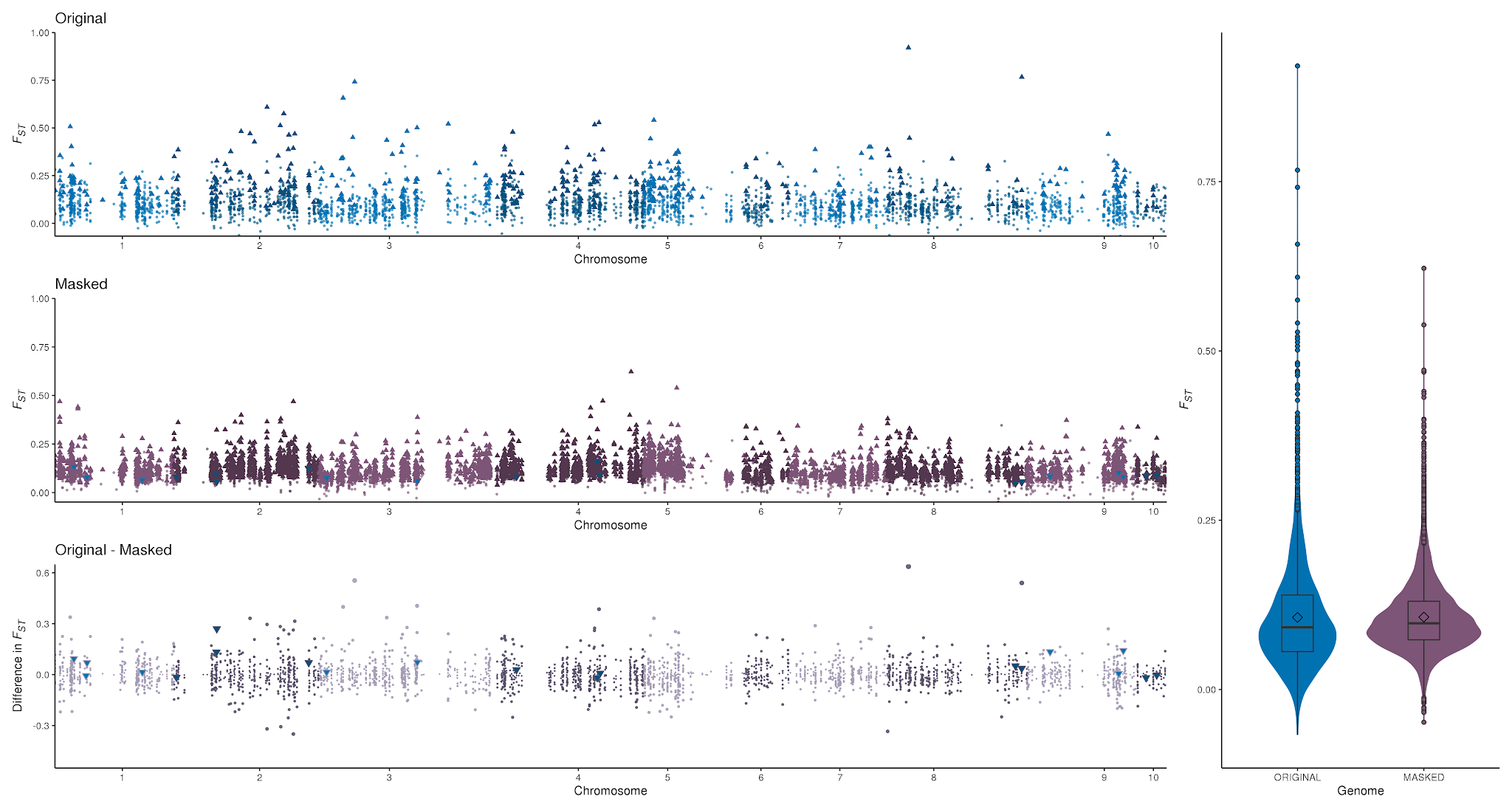
<https://github.com/The-Eastern-Oyster-Genome-Project/2022_Eastern_Oyster_Haplotig_Masked_Genome/raw/main/Comparative_Analysis/Output/Figures/Supplemental/Figure.S5.Full.FST.diplo.png>

**Figure S6- Comparison of estimates of *F_ST_* across new diplotigous regions in the original and haplotig-masked genomes.** Panel (A) is values of *F_ST_* averaged across 10 kb windows in the original genome that changed to diplotigous coverage levels after haplotig-masking. Windows that contained any outlier SNP loci were changed to triangles (483 windows). Panel (B) panel is values of *F_ST_* averaged across in the same 10 kb newly diplogtigous windows across the haplotig-masked genome. Windows that contained any outlier SNP loci were changed to triangles (4,527 windows) . There were 20 windows with SNPs that were identified as outliers in the original genome analysis but not in the haplotig-masked analysis, and these windows are marked as upside down blue triangles. Panel (C) is the difference between the original and the haplotig-masked values in 10 kb windows across the entire genome with dot size directly related to the distance from zero. There were 20 windows with SNPs that were identified as outliers in the original genome analysis but not in the haplotig-masked analysis, and these windows are marked as upside down blue triangles. Panel (D) is a violin and boxplot of 10kb averaged values between the two genome versions. For all plots, blue is the original genome and purple is the haplotig-masked genome.

### Figure S7- Comparison of estimates of FST across newly diplotigous regions using the low structure subset (LSS)

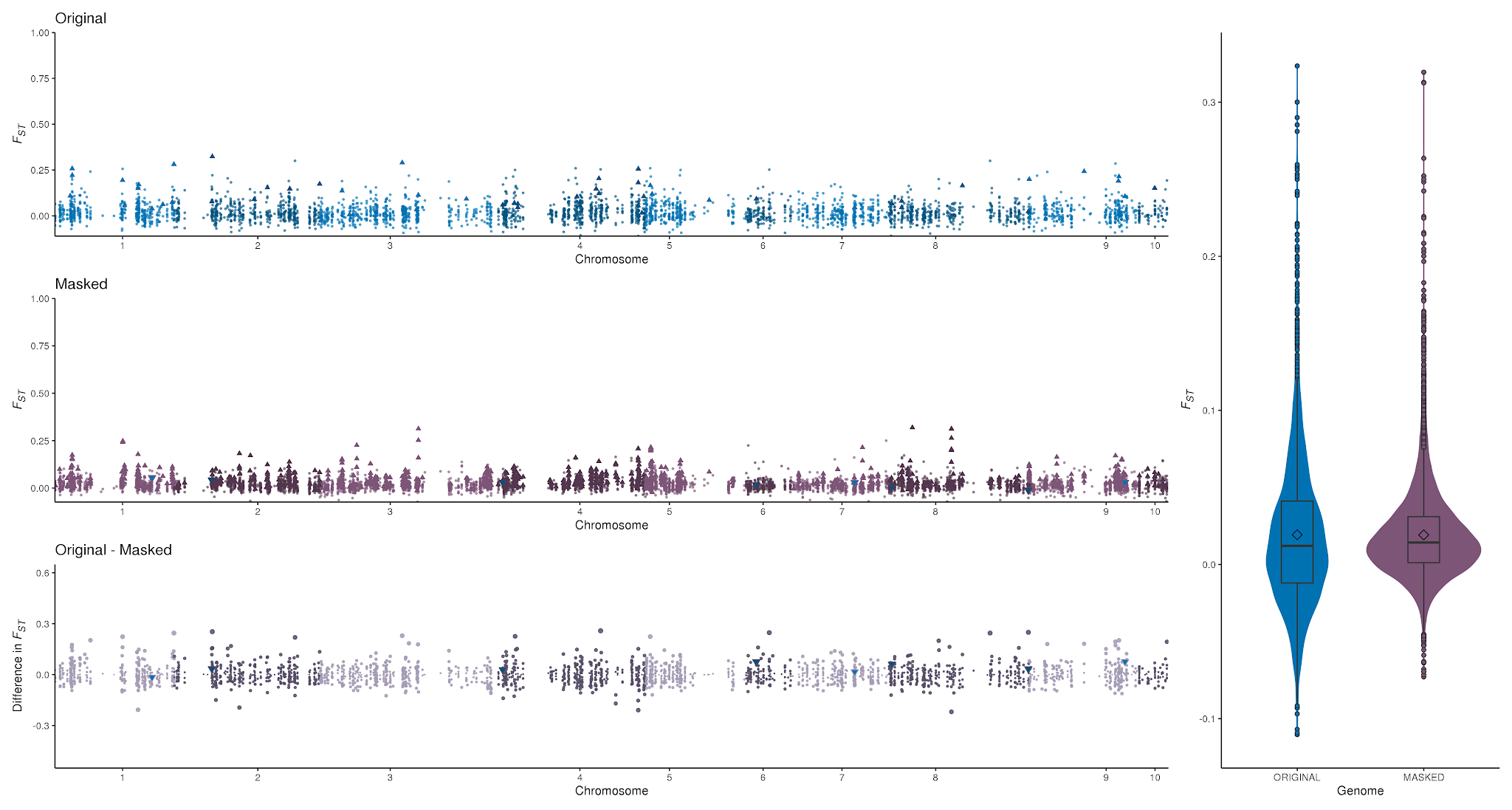
<https://github.com/The-Eastern-Oyster-Genome-Project/2022_Eastern_Oyster_Haplotig_Masked_Genome/raw/main/Comparative_Analysis/Output/Figures/Supplemental/Figure.S6.LSS.FST.diplotigs.png>

**Figure S7- Comparison of estimates of *F_ST_* across newly diplotigous regions using the low structure subset (LSS).** Estimates of *F_ST_* were calculated using the low structure subset (LSS) to examine how haplotigs affect population structure inference in a lower signal system. Panel (A) is values of *F_ST_* averaged across 10 kb windows in the original genome that changed to diplotigous coverage levels after haplotig-masking. Windows that contained any outlier SNP loci were changed to triangles (55 windows). Panel (B) panel is values of *F_ST_* averaged across in the same 10 kb newly diplogtigous windows across the haplotig-masked genome. Windows that contained any outlier SNP loci were changed to triangles (860 windows) . There were 8 windows with SNPs that were identified as outliers in the original genome analysis but not in the haplotig-masked analysis, and these windows are marked as upside down blue triangles. Panel (C) is the difference between the original and the haplotig-masked values in 10 kb windows across the entire genome with dot size directly related to the distance from zero. There were 8 windows with SNPs that were identified as outliers in the original genome analysis but not in the haplotig-masked analysis, and these windows are marked as upside down blue triangles. Panel (D) is a violin and boxplot of 10kb averaged values between the two genome versions. For all plots, blue is the original genome and purple is the haplotig-masked genome.

### Figure S8- Comparison of estimates of *V_ST_* across the original and haplotig-masked genomes

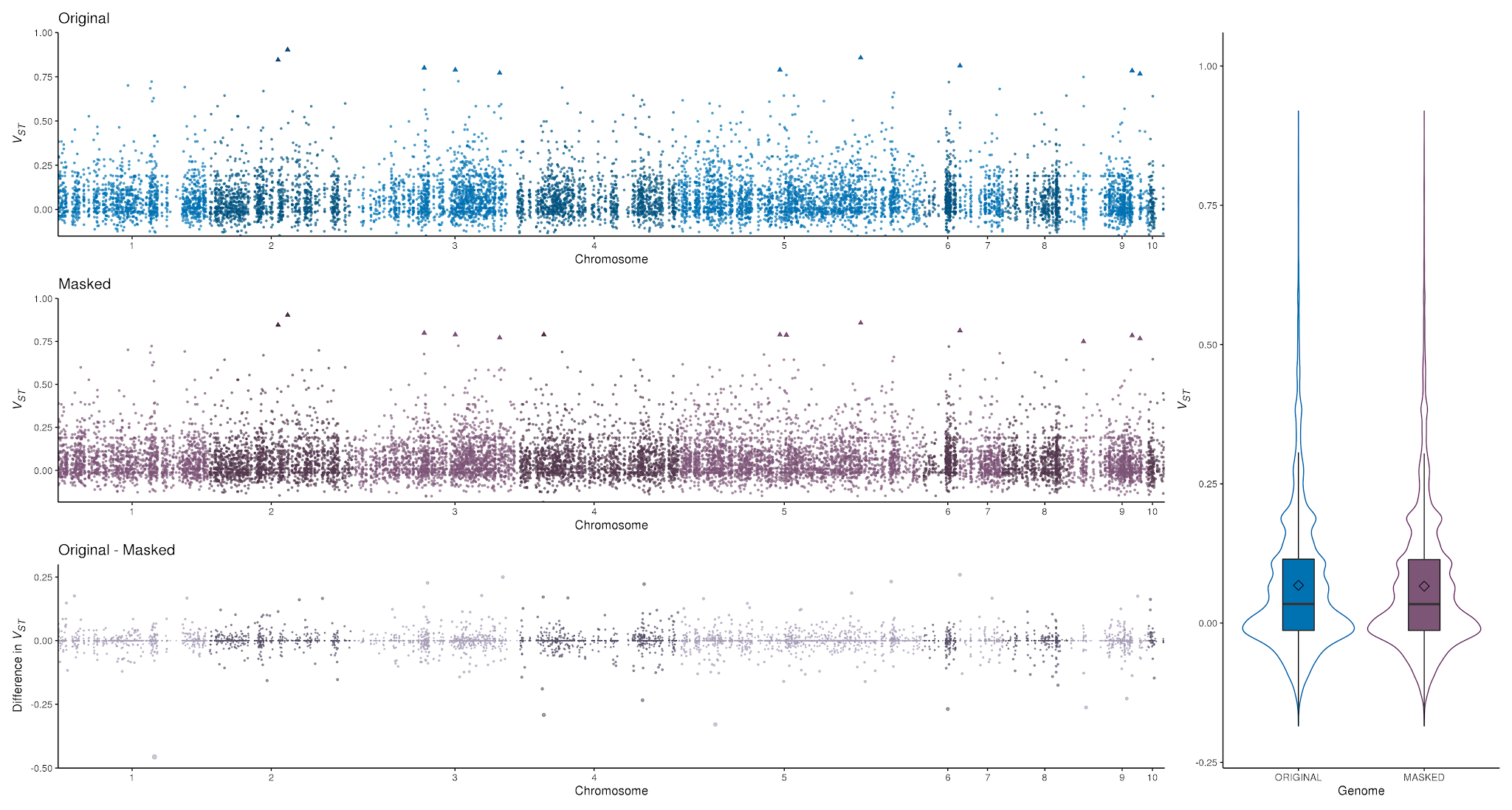
<https://github.com/The-Eastern-Oyster-Genome-Project/2022_Eastern_Oyster_Haplotig_Masked_Genome/raw/main/Comparative_Analysis/Output/Figures/Supplemental/Figure.S7.all.VSTs.man.png>

**Figure S8- Comparison of estimates of *V_ST_* across the original and haplotig-masked genomes.** Panel (A) is values of *V_ST_* averaged across 10 kb windows in the original genome. Panel (B) panel is values of *V_ST_* averaged across 10 kb windows across the haplotig-masked genome. Panel (C) is the difference between the original and the haplotig-masked values in 10 kb windows across the entire genome with dot size directly related to the distance from zero. Panel (D) is a violin and boxplot of 10kb averaged values between the two genome versions. For all plots, blue is the original genome and purple is the haplotig-masked genome.

### Figure S9- Comparison of estimates of *V_ST_* across the original and haplotig-masked genomes using the low structure subset (LSS)

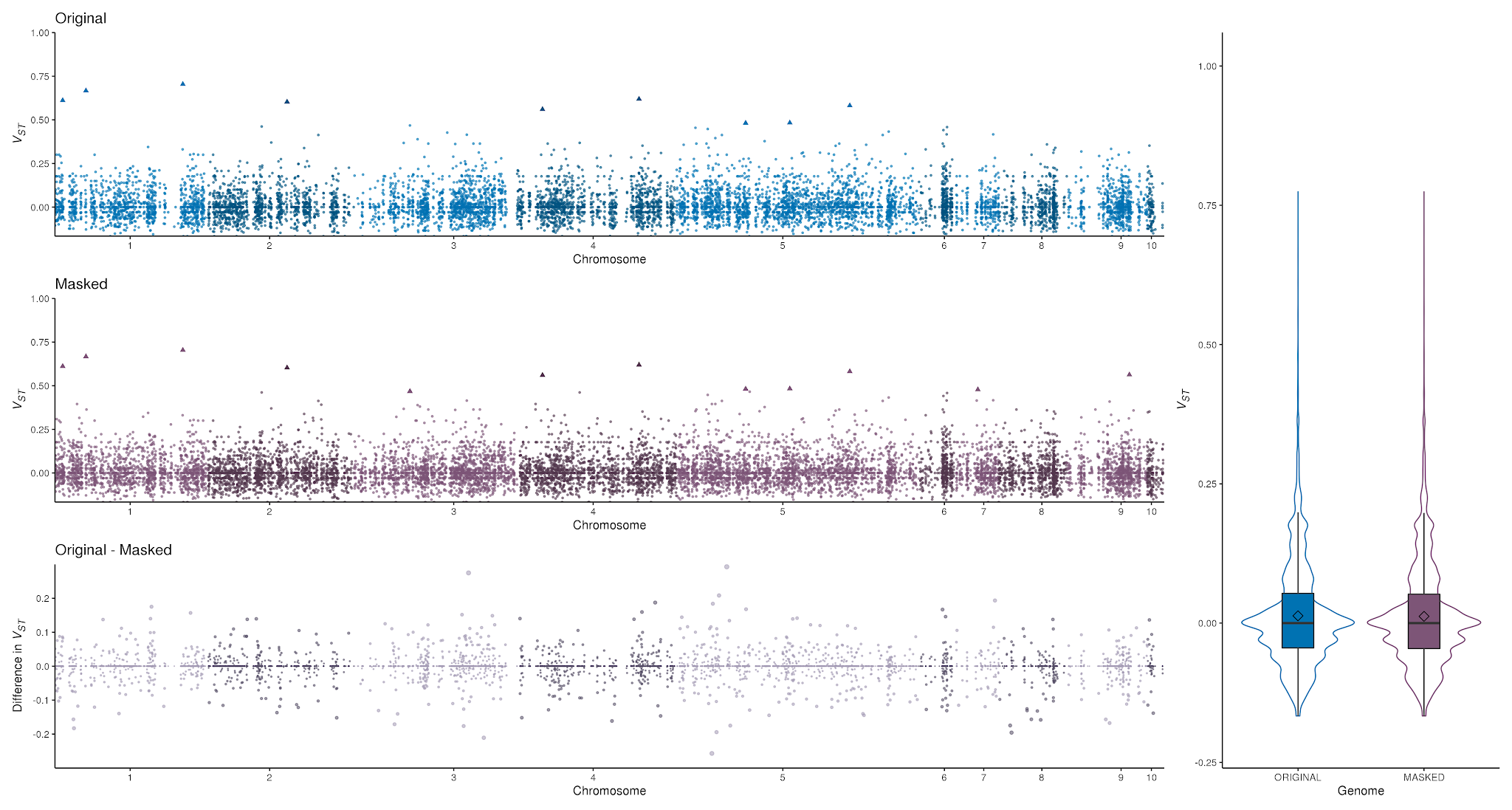

**Figure S9- Comparison of estimates of *V_ST_* across the original and haplotig-masked genomes using the low structure subset (LSS).** Estimates of *V_ST_* were calculated using the low structure subset (LSS) to examine how haplotigs affect population structure inference in a lower signal system. Panel (A) is values of *V_ST_* averaged across 10 kb windows in the original genome. Panel (B) panel is values of *V_ST_* averaged across 10 kb windows across the haplotig-masked genome. Panel (C) is the difference between the original and the haplotig-masked values in 10 kb windows across the entire genome with dot size directly related to the distance from zero. Panel (D) is a violin and boxplot of 10kb averaged values between the two genome versions. For all plots, blue is the original genome and purple is the haplotig-masked genome.
